# Large palindromes on the primate X Chromosome are preserved by natural selection

**DOI:** 10.1101/2020.12.29.424738

**Authors:** Emily K. Jackson, Daniel W. Bellott, Ting-Jan Cho, Helen Skaletsky, Jennifer F. Hughes, Tatyana Pyntikova, David C. Page

## Abstract

Mammalian sex chromosomes carry large palindromes that harbor protein-coding gene families with testis-biased expression. However, there are few known examples of sex-chromosome palindromes conserved between species. We identified 26 palindromes on the human X Chromosome, constituting more than 2% of its sequence, and characterized orthologous palindromes in the chimpanzee and the rhesus macaque using a clone-based sequencing approach that incorporates full-length nanopore reads. Many of these palindromes are missing or misassembled in the current reference assemblies of these species’ genomes. We find that 12 human X palindromes have been conserved for at least 25 million years, with orthologs in both chimpanzee and rhesus macaque. Insertions and deletions between species are significantly depleted within the X palindromes’ protein-coding genes compared to their non-coding sequence, demonstrating that natural selection has preserved these gene families. Unexpectedly, the spacers that separate the left and right arms of palindromes are a site of localized structural instability, with 7 of 12 conserved palindromes showing no spacer orthology between human and rhesus macaque. Analysis of the 1000 Genomes Project dataset revealed that human X-palindrome spacers are enriched for deletions relative to arms and flanking sequence, including a common spacer deletion that affects 13% of human X Chromosomes. This work reveals an abundance of conserved palindromes on primate X Chromosomes, and suggests that protein-coding gene families in palindromes (most of which remain poorly characterized) promote X-palindrome survival in the face of ongoing structural instability.

## INTRODUCTION

The human X Chromosome contains two classes of genomic sequence with distinct evolutionary histories (Mueller et al. 2013). Most human X-Chromosome sequence is ancestral, containing genes retained from the ancestral autosome from which the mammalian sex chromosomes evolved some 200-300 million years ago (Ohno 1967, Lahn and Page 1999); these ancestral genes display diverse patterns of expression. However, around 2% of the human X Chromosome comprises ampliconic sequence, which contains gene families that are expressed predominantly in testis and that were not present on the ancestral autosome (Mueller et al. 2013). Amplicons are long, highly identical repeat units, with lengths that can exceed 100 kb in length and sequence identities >99% (Kuroda-Kawaguchi et al. 2001). Although some amplicons form tandem arrays, the majority comprise large palindromes, with mirrored repeats facing head to head (Skaletsky et al. 2003). The testis-biased expression of human X-Chromosome palindrome gene families has led to speculation that X palindromes play roles in spermatogenesis (Warburton et al. 2004, Mueller et al. 2013), consistent with evolutionary theories predicting that the X chromosome should preferentially accumulate male-beneficial alleles (Rice 1984). In contrast to ancestral sequence, however, few studies have focused on X-Chromosome palindromes, and X palindromes have not been characterized in non-human primates. As a result, little is known about the origins of X-Chromosome palindromes or the evolutionary forces that shape them.

Studies of X-Chromosome palindromes have been impeded by the difficulty of obtaining accurate reference sequence. Segmental duplications are commonly collapsed by assembly algorithms into a single repeat unit (Eichler 2001), and they are severely underrepresented in reference genomes assembled using short sequencing reads and whole-genome shotgun (WGS) algorithms (Alkan et al. 2011). Several mammalian Y Chromosomes, which also contain palindromes and other amplicons (Skaletsky et al. 2003, Hughes et al. 2010, Hughes et al. 2012, Soh et al. 2014, Hughes et al. 2020), were sequenced using labor-intensive but highly accurate clone-based approaches (Kawaguchi-Kuroda et al. 2001, Bellott et al. 2018). Recently, the incorporation of ultralong nanopore reads into a clone-based sequencing approach (Single Haplotype Iterative Mapping and Sequencing 3.0, or SHIMS 3.0) has enabled the time-and cost-effective resolution of amplicons, including the *TSPY* array on the human Y Chromosome, that had been impervious to previous assembly methods (Bellott et al. 2020). Accurate representation of amplicons and other segmental duplications in mammalian genomes is particularly important given the disproportionate roles of segmental duplications in mediating deletions, duplications, inversions, and other complex rearrangements across the human genome (Stankiewicz and Lupski 2010).

Only two mammalian X Chromosomes have been sequenced using a high-resolution, clone-based approach: the mouse X Chromosome (Church et al. 2009) and the human X Chromosome (Ross et al. 2005, Mueller et al. 2013). Palindromes containing testis-biased gene families are abundant on both the human and mouse X Chromosomes (Warburton et al. 2004, Mueller et al. 2008); however, palindrome gene content largely does not overlap between the two species, suggesting that the palindromes are not orthologous (Mueller et al. 2013). A subset of human X-palindrome gene families have highly divergent orthologs on the chimpanzee X Chromosome, but without a complete reference sequence, their copy number and orientations could not be determined (Stevenson et al. 2007). To determine whether human X palindromes have orthologs in other primates, we used SHIMS 3.0 to generate high-quality reference sequence for portions of the X Chromosome that are orthologous to human X palindromes in two non-human primate species, the chimpanzee and the rhesus macaque.

## RESULTS

### Characterization of 26 palindromes on the human X Chromosome

To ensure consistency in palindrome annotation between species, we began by re-annotating human X palindromes. We identified palindromic amplicons by searching the reference sequence of the human X Chromosome for inverted repeats > 8 kb in length (which excludes repetitive elements such as LINEs and SINEs) that display >99% nucleotide identity (Fig. 1A). We used a kmer-based method (Teitz et al 2018) to precisely define the coordinates of each palindrome (see Methods), and triangular dot plots to visualize palindromes and other genomic repeats (Fig. 1B, C). In total, we identified 26 palindromic amplicons on the human X Chromosome (Table 1, Supplemental Table S1); these palindromes, including both arms and spacer sequence, comprise over 3.46 Mb, or 2.2% of the length of the human X Chromosome. Palindrome arm lengths range from 8 kb to 140 kb, with arm-to-arm identities as high as 99.99%, representing one nucleotide difference between arms per 10,000 base pairs. Palindrome spacer lengths span several orders of magnitude, ranging from 77 bp to 358 kb. The vast majority of palindromes (21 of 26) have at least one protein-coding gene in their arms. No primate Y palindromes have been observed with protein-coding genes in their spacers (Skaletsky et al. 2003, Hughes et al. 2010, Hughes et al. 2012), but we found that 11 of 26 human X palindromes also contain at least one protein-coding gene in their spacer.

**Figure 1.**
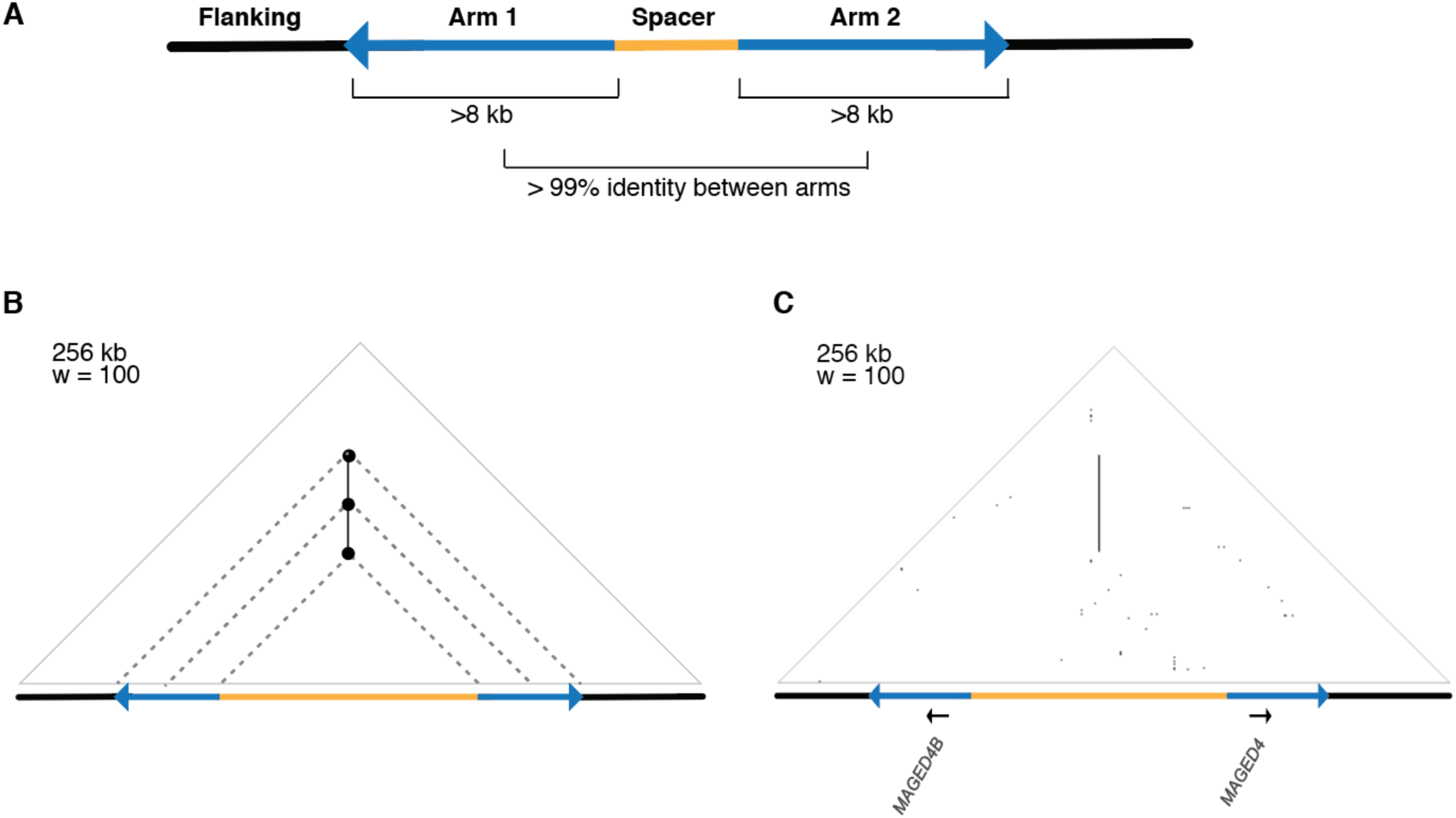
Overview of human X-Chromosome palindromes. a) Schematic of a palindrome. b) Schematic of a triangular dot plot. Dots are placed at a 90-degree angle between identical kmers, or “words,” within a DNA sequence. Palindromes appear as vertical lines. “w”: word size used to construct the dot plot. c) Triangular dot plot for human X palindrome P3, including annotated protein-coding genes.

**Table 1.**
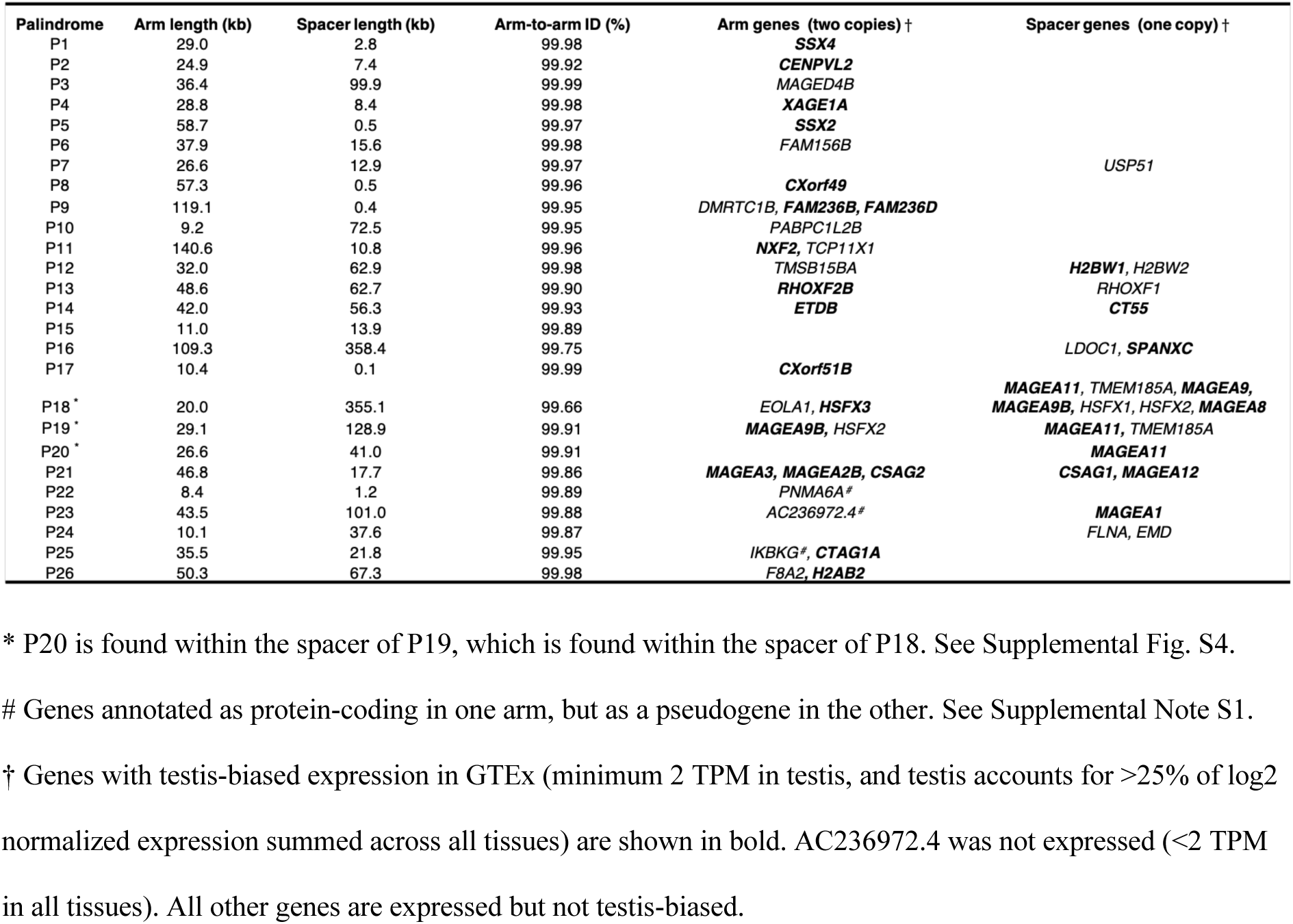
Palindromes on the human X Chromosome

To investigate the functions of palindrome-encoded genes, we examined the expression patterns of all 45 human X-palindrome arm and spacer genes using data from the Genotype-Tissue Expression (GTEx) Consortium (The GTEx Consortium 2017). Palindrome arm genes are present in two nearly identical copies, and RNA sequencing reads map equally well to both, which poses a problem for accurately estimating the expression level of genes in palindrome arms using traditional software that discards multi-mapping reads, as was recently shown for ampliconic genes on the Y Chromosome (Godfrey et al 2020). We therefore re-analyzed GTEx data with kallisto, which assigns multi-mapping reads based on a probability distribution (Bray et al. 2016, Godfrey et al. 2020). We found that 18 of 30 gene families (60%) in palindrome arms were expressed predominantly in testis, consistent with previous reports (Table 1, Supplemental Fig. S1) (Warburton et al. 2004). Genes in palindrome spacers showed similar expression patterns: 10 of 17 genes (58.8%) were expressed predominantly in testis (Table 1, Supplemental Fig. S1), suggesting that reproductive specialization of palindrome genes includes both arm genes and spacer genes. We further examined the expression of testis-biased gene families in palindrome arms and spacers across human spermatogenesis using bulk RNA-Seq data from Jan et al. 2017 (Supplemental Fig. S2). Across the 20 testis-biased gene families with detectable expression at one or more timepoints, we observed patterns ranging from highest expression in spermatogonia to highest expression in round spermatids, suggesting that human X palindrome gene families play roles across multiple stages of spermatogenesis.

Palindromes on the human Y chromosome are depleted for LINEs and other transposable elements (Skaletsky et al. 2003). We used RepeatMasker to examine the density of transposable elements in human X palindrome arms, and found that transposable elements (LINEs, SINEs, LTRs, and DNA elements) account for 50.2% of palindrome arms, compared to 58.1% of flanking sequence (p<0.05, Mann-Whitney U). The density of transposable elements is negatively correlated with recombination rates across mammalian genomes, which may reflect the increased efficiency of natural selection in removing mildly deleterious insertions (Jensen-Seaman et al. 2004). We conclude that transposable elements are depleted both in X and Y palindromes in humans, and that this likely results from elevated recombination rates in palindrome arms (Rozen et al. 2003).

### Generation of high-resolution X-palindrome reference sequence for chimpanzee and rhesus macaque

To understand the origins of human X palindromes, we searched for orthologous palindromes in two non-human primates, the chimpanzee and the rhesus macaque. Existing reference genomes for chimpanzee and rhesus were not generated using a clone-based approach comparable to that for the human genome, so to address this limitation, we generated 14.43 Mb of non-redundant high-quality reference sequence for portions of the chimpanzee and rhesus X Chromosomes that are orthologous to human palindromes (Fig. 2A). We used the recently developed SHIMS 3.0 method, a clone-based approach that incorporates both Illumina and nanopore reads (Bellott et al. 2020). This method provides high structural confidence due to the generation of full-length nanopore reads spanning each clone: For 83 of 107 clones sequenced using SHIMS 3.0, we were able to verify the accuracy of internal repeat structures by comparing the finished sequence to one or more full-length nanopore reads (Fig. 2B, Supplemental Table S2).

**Figure 2.**
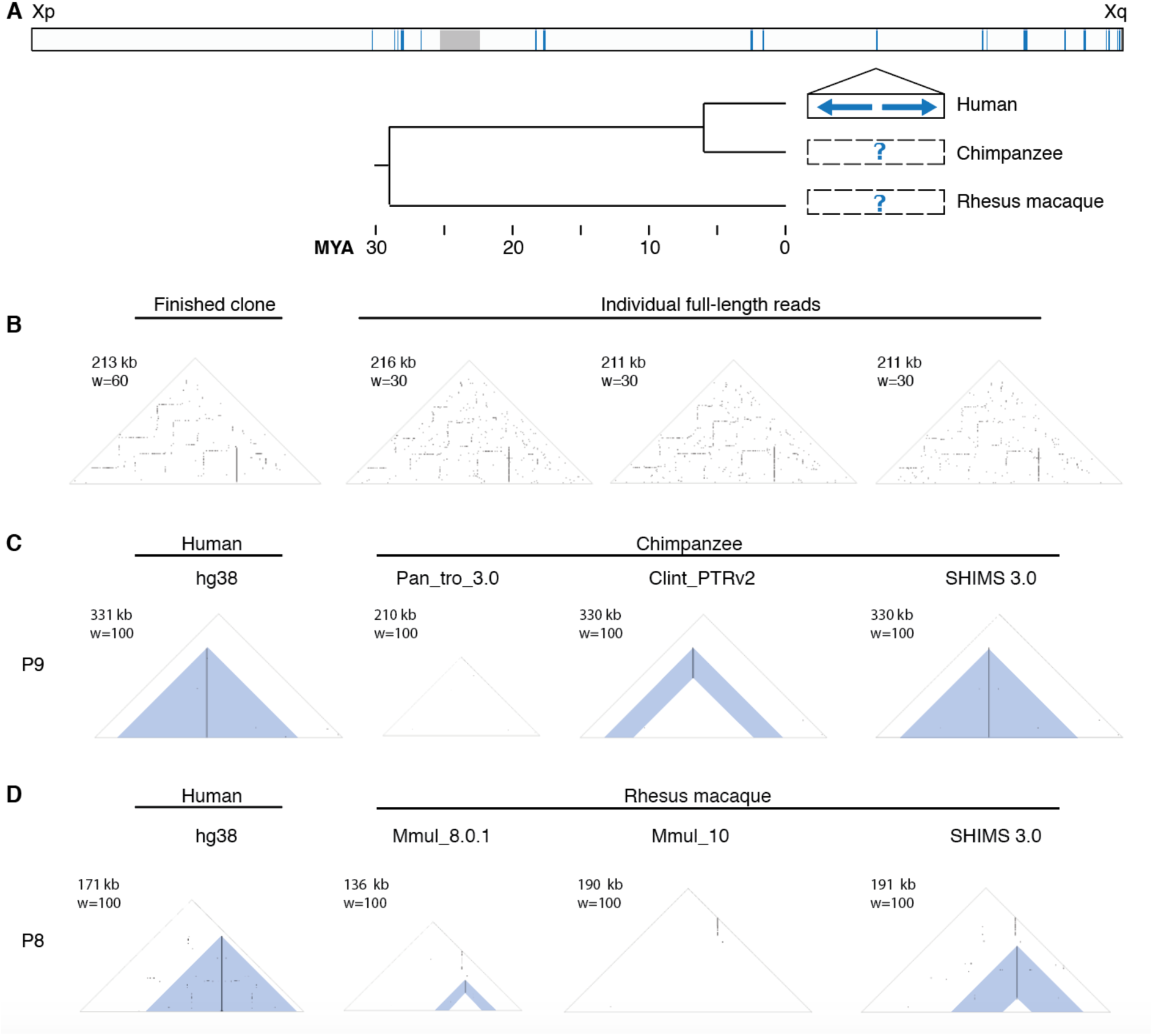
Improvements to prior reference assemblies for chimpanzee and rhesus macaque. a) Sequencing approach. Top bar: Locations of 26 human X palindromes (blue bands). Gray band shows centromere location. Below: Expansion of a single region containing a human X palindrome (solid black box). One or more clones were selected to span orthologous regions in chimpanzee and rhesus macaque (dashed black boxes). Tree shows estimated divergence times from TimeTree (Kumar et al. 2017). b) Full-length nanopore reads supporting the structure of a single finished chimpanzee clone (CH251-385I8). c,d) Two primate X palindromes resolved using SHIMS 3.0 that were missing or misassembled in existing X-Chromosome assemblies. c) Triangular dot plots from the region orthologous to human P9 in chimpanzee assemblies Pan_tro_3.0 (Kuderna et al. 2017), Clint_PTRv2 (Kronenberg et al. 2018), and SHIMS 3.0. d) Triangular dot plots from the region orthologous to human P8 in rhesus macaque assemblies Mmul_8.0.1 (Zimin et al. 2014), Mmul_10 (Warren et al. 2020), and SHIMS 3.0.

Our assemblies revealed 39 palindromes in total on the chimpanzee and rhesus macaque X Chromosomes, collectively comprising 4.90 Mb of palindromic amplicon sequence. Only 10 of these palindromes (25.6%) were represented accurately in existing X-Chromosome assemblies that were generated using primarily short-read whole genome shotgun (WGS) approaches (chimpanzee, Pan_tro_3.0; rhesus macaque, Mmul_8.0.1) (Fig. 2C,D; Supplemental Fig. S3; Supplemental Tables S3,S4) (Kuderna et al. 2017, Zimin et al. 2014). We also compared our SHIMS 3.0 assemblies to two Figure 2. Improvements to prior reference assemblies for chimpanzee and rhesus macaque. a) Sequencing approach. Top bar: Locations of 26 human X palindromes (blue bands). Gray band shows centromere location. Below: Expansion of a single region containing a human X palindrome (solid black box). One or more clones were selected to span orthologous regions in chimpanzee and rhesus macaque (dashed black boxes). Tree shows estimated divergence times from TimeTree (Kumar et al. 2017). b) Full-length nanopore reads supporting the structure of a single finished chimpanzee clone (CH251-385I8). c,d) Two primate X palindromes resolved using SHIMS 3.0 that were missing or misassembled in existing X-Chromosome assemblies. c) Triangular dot plots from the region orthologous to human P9 in chimpanzee assemblies Pan_tro_3.0 (Kuderna et al. 2017), Clint_PTRv2 (Kronenberg et al. 2018), and SHIMS 3.0. d) Triangular dot plots from the region orthologous to human P8 in rhesus macaque assemblies Mmul_8.0.1 (Zimin et al. 2014), Mmul_10 (Warren et al. 2020), and SHIMS 3.0. assemblies generated using long-read WGS approaches incorporating PacBio reads (chimpanzee, Clint_PTRv2; rhesus macaque, Mmul_10) (Kronenberg et al. 2018, Warren et al. 2020). Here, we found that while 18 palindromes (46.2%) were represented accurately, 21 palindromes (53.8%) remained missing or incomplete (Fig. 2C,D; Supplemental Fig. S3; Supplemental Tables S3, S4). Palindromes that were missing or incomplete in existing long-read WGS assemblies had longer arms than palindromes that were represented accurately (Clint_PTRv2: 63 kb versus 20 kb; Mmul_10: 65 kb versus 22 kb) (p<0.01 for both comparisons, Mann-Whitney U), suggesting that large palindromes remain particularly intractable to whole-genome shotgun approaches (Supplemental Tables S3, S4). We therefore carried out all subsequent analyses using our newly generated SHIMS 3.0 assemblies.

### Conservation of X palindromes in two non-human primates

We annotated palindromes in chimpanzee and rhesus macaque using the same criteria used to annotate human palindromes (minimum 8-kb arm length, and minimum 99% nucleotide identity between arms). We also included one palindrome in rhesus macaque with arms of 6.5 kb, given that the palindrome exhibited 99% arm-to-arm identity and was orthologous to human palindrome P10 (Supplemental Fig. S5). In total, we discovered 21 palindromes in chimpanzee and 18 palindromes in rhesus macaque, demonstrating that X-linked palindromes are not unique to humans but represent a common feature of primate X Chromosomes, at least to Old World monkeys (Fig. 3A).

**Figure 3.**
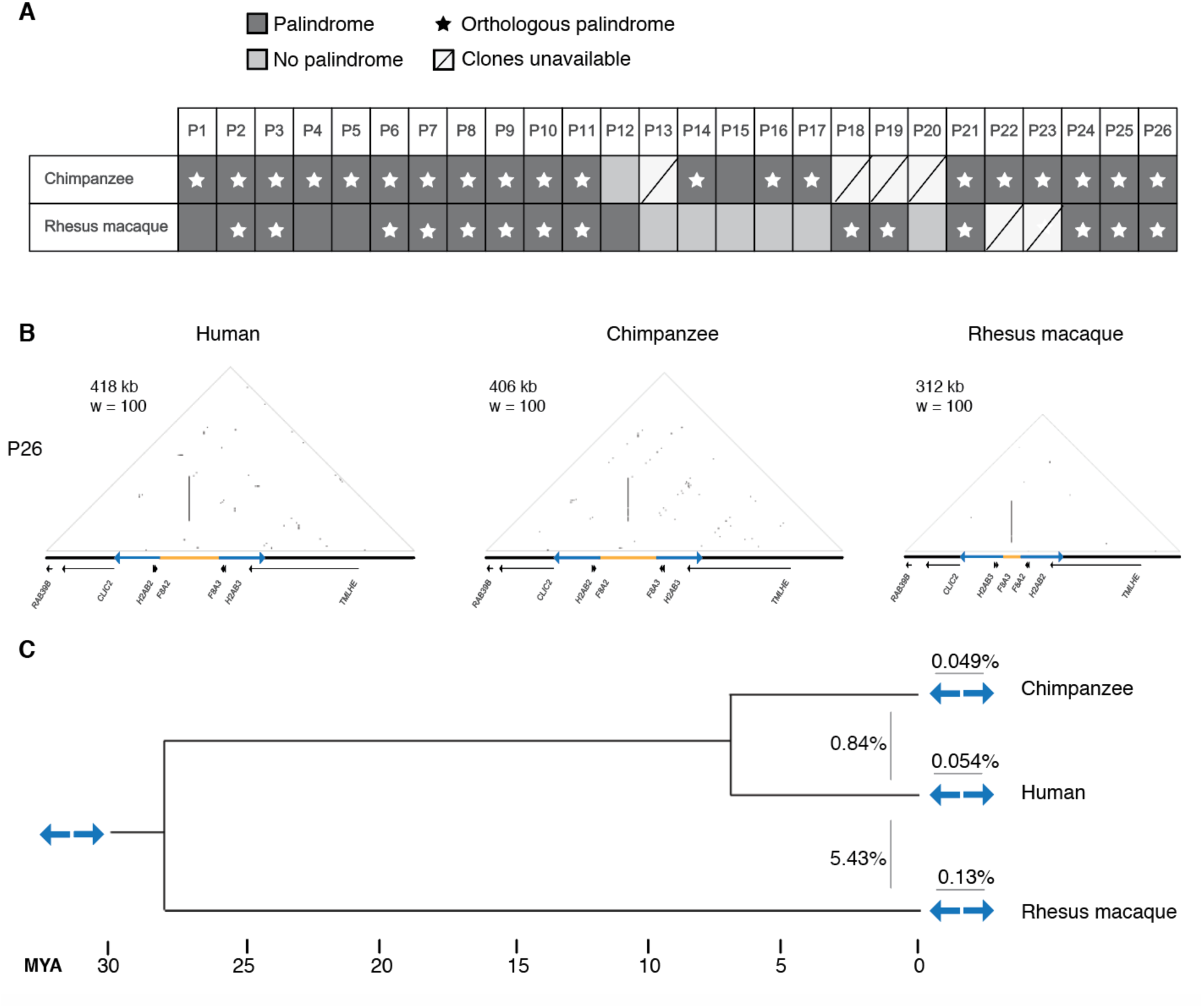
Conservation of X-Chromosome palindromes across primates. a) Conservation status of 26 human palindromes in chimpanzee and rhesus macaque. b) Triangular dot plots for a palindrome (P26) conserved between human, chimpanzee, and rhesus macaque. Images are to scale. c) Arm-to-arm divergence within species (above blue arrows) versus between species (left of blue arrows). Values are the average percent divergence across 12 palindromes shared by human, chimpanzee, and macaque. Divergence times estimated using TimeTree (Kumar et al. 2017).

We applied two criteria to identify orthologs of human X palindromes (Supplemental Fig. S4A, B). First, we required that at least 20% of each palindrome arm in chimpanzee or rhesus macaque align to its putative human ortholog. This excluded three palindromes; we suggest that despite their similar genomic positions, these palindromes arose independently in each lineage (Supplemental Fig. S4C). Second, we required that the portion of palindrome arms that aligned between species have minimal alignment to flanking sequence, defined through high-quality BLAST hits (see Methods). This excluded two palindromes in which the palindrome arm in rhesus macaque mapped equally well to more than two locations in the human sequence, including flanking sequence, consistent with scenarios in which similar palindromes arose independently in each species (Supplemental Fig. S4D). We note that this approach for defining orthologous palindromes is conservative, and may exclude palindromes that were present in the common ancestor but underwent extreme rearrangements in one or more species.

After excluding four regions for which we were not able to generate SHIMS 3.0 assemblies, we found that the vast majority of the remaining human palindromes—20 of 22—have an orthologous palindrome in chimpanzee; the same is true for 14 of 24 palindromes in rhesus macaque (Fig. 3A). For each species, we annotated protein-coding genes in our newly generated sequence, and constructed dot plots with accompanying tracks showing the positions and orientations of palindrome arms and genes for each region (Fig. 3B, Supplemental Fig. S5). Comparisons of human sequence with non-human primate sequence demonstrated that these palindromes harbor orthologous protein-coding genes (Fig. 3B). Previous literature has reported that nucleotide identity between sex chromosome palindrome arms is maintained by ongoing gene conversion (Skaletsky et al. 2003, Rozen et al. 2003, Swanepoel et al. 2020); consistent with this, we find that the average arm-to-arm divergence within a species (<0.2% for all species) is lower than the average arm-to-arm divergence between species (0.87% and 5.50% for human versus chimpanzee and human versus rhesus macaque, respectively) (Fig. 3C). We conclude that nearly half of human X palindromes — 12 of 26 — were present in the common ancestor of human, chimpanzee, and rhesus macaque at least 25 million years ago, and have been maintained by ongoing gene conversion in all three lineages since their divergence. We subjected these 12 palindromes shared by human, chimpanzee, and macaque to further analyses to uncover the processes governing their evolution and unexpectedly deep conservation.

### Structural changes between species are concentrated around the center of symmetry

Palindromes on the human X and Y Chromosomes are associated with a wide range of pathogenic rearrangements, the majority of which result from non-allelic homologous recombination (NAHR) between near-identical palindrome arms (Lakich et al. 1993, Small et al. 1997, Aradhya et al. 2001, Lange et al. 2009, Scott et al. 2010). Although palindromes are well-known sites of genomic instability in humans, little is known about the stability of palindromic structures on longer evolutionary timescales. We used our set of 12 X palindromes shared by human, chimpanzee, and macaque to investigate structural changes in the palindromes over time.

NAHR between palindrome arms has two common outcomes: NAHR within a single chromatid results in inversions of the spacer, while NAHR between misaligned sister chromatids results in acentric and dicentric fragments (Lange et al. 2009). Given that acentric and dicentric fragments are not expected to be stably inherited, we instead looked for evidence of spacer inversions between species. Inversions within X palindromes have been previously reported to exist as neutral polymorphisms in human populations (Small et al. 1997), as well as pathogenic rearrangements (Lakich et al. 1993, Porubsky et al. 2020). We found abundant evidence for inversions in primate X palindromes: In cases where the orientation of the spacer could be confidently determined, the frequency of inversions between chimpanzee and human, and between human and rhesus macaque, was about 50% (Fig. 4A, Supplemental Fig. S5). This is consistent with the notion that inversions are common events, the majority of which are not harmful, except in rare instances where they disrupt a protein-coding gene. Our SHIMS 3.0 assemblies each derive from a single male individual, but in light of the results from Small and colleagues, who found that human P24 spacers are inverted in 18% of European X Chromosomes, we would anticipate that inversion polymorphisms exist within all three species. Indeed, a recent study found that 10 out of 13 newly identified X chromosome inversion polymorphisms in chimpanzee and/or bonobo occurred in X palindromes (Porubsky et al. 2020).

**Figure 4.**
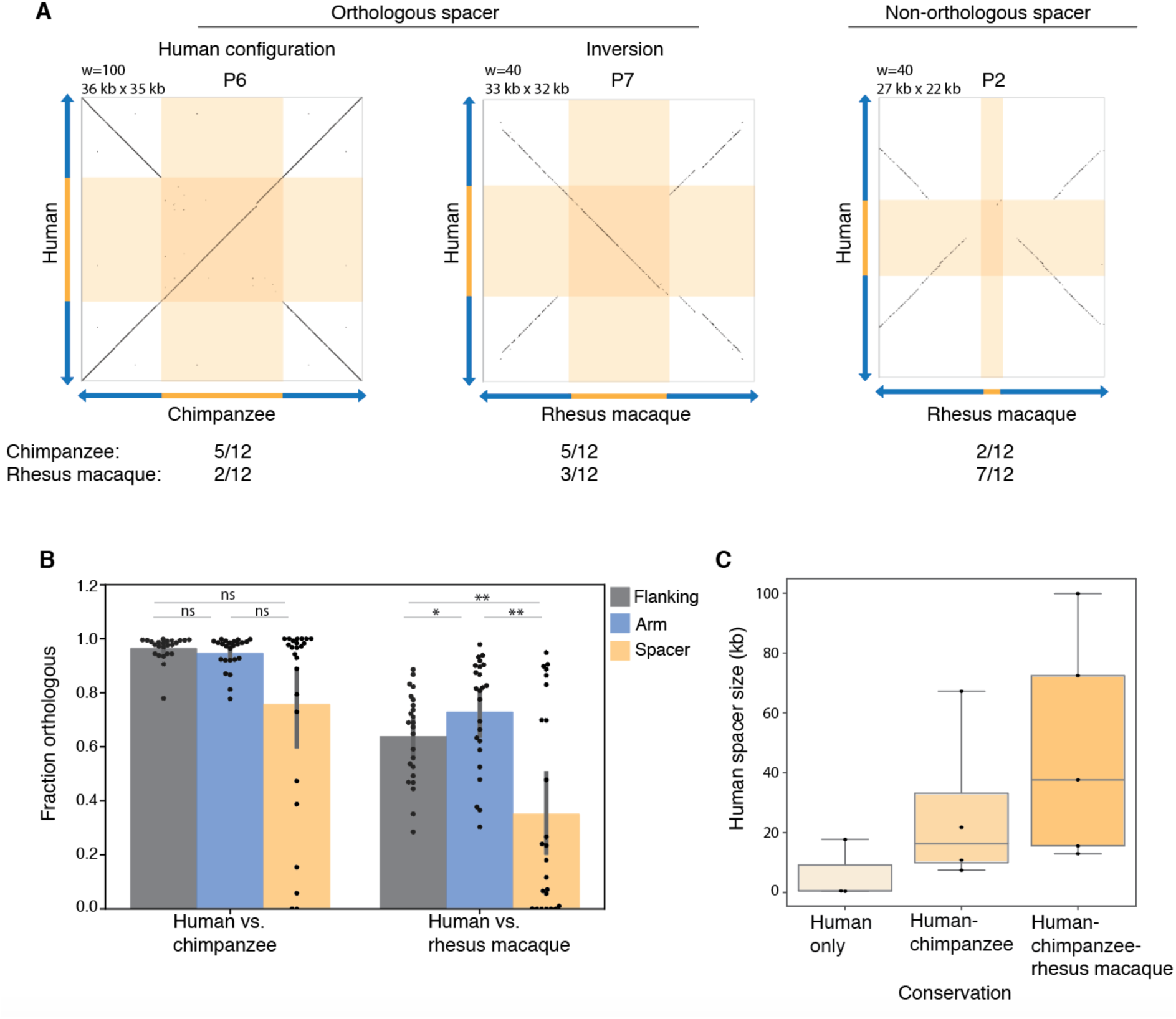
Structural changes between orthologous X-chromosome palindromes are concentrated around the center of symmetry. a) Square dot plots comparing the center of the palindrome, including the spacer and 10 kb of inner arm sequence on each side, between the indicated species. “Orthologous spacer”: > 20% of the spacer from one species aligned to the spacer from the other, in either the same orientation (“Human configuration”) or opposite orientation (“Inversion”). “Non-orthologous”: < 20% of the spacer from one species aligned to the spacer from the other. Values show the number of orthologous palindromes shared by human, chimpanzee, and macaque in each category. b) Average fraction of sequence that could be aligned between species. *p < 0.05, **p <0.01, Mann-Whitney U. c) Sizes of human spacers, binned according to the species between which they are conserved.

Unexpectedly, we observed numerous examples in which spacer sequence could not be aligned between species, despite robust alignment of most or all of the palindrome arms (Fig. 4A, Supplemental Fig. S6). This phenomenon was observed in 2 of 12 spacer comparisons between human and chimpanzee, and 7 of 12 spacer comparisons between human and rhesus macaque (Fig. 4A). Non-orthologous spacers could be the result of insertions, deletions, or translocations of sequence within a palindrome. Rather than attempting to reconstruct each event, we used the fraction of sequence that is orthologous as a simplified metric for sequence rearrangements, and asked whether rearrangements are concentrated within spacers compared to palindrome arms or flanking sequence. We aligned each spacer between species, calculated the fraction of orthologous sequence, and repeated this for palindrome arms and flanking sequence. When comparing human versus chimpanzee, the average fraction of orthologous sequence was lowest in spacers (75.8% for spacers versus 96.5% and 94.7% for flanking sequence and arms, respectively), though the result did not reach statistical significance (Fig. 4B). When comparing human versus rhesus macaque, the average fraction of orthologous sequence was significantly lower in spacers (35.2%) than in arms (72.9%) or flanking sequences (63.8%) (p<0.01 for both, Mann-Whitney U) (Fig. 4B). Consistent with our observations in Fig. 4A, these results were driven by a subset of palindromes in which the spacer displayed little or no similarity between species, rather than a small and consistent decrease in similarity affecting all palindromes equally (Fig. 4B). Spacer size was positively correlated with the degree of conservation between species, suggesting that small spacers may be particularly unstable (Fig. 4C). We conclude that in addition to inversions, palindromes rearrangements are concentrated around the center of symmetry, and that palindromes with small spacers are most susceptible to rearrangement.

### Natural selection has preserved palindrome gene families

We wondered whether natural selection might provide a countervailing force against structural instability in primate X palindromes. Among the 12 human X palindromes with orthologous palindromes in chimpanzee and rhesus macaque, there are 17 protein-coding gene families in palindrome arms, and 5 protein-coding genes in palindrome spacers (Supplemental Table S5). The functions of these gene families are poorly characterized in humans: Only 1 of 17 human palindrome arm gene families (6%) have phenotypes listed in the Online Mendelian Inheritance in Man (OMIM) database (McKusick-Nathans Institute of Genetic Medicine 2020). This represents a four-fold depletion relative to other protein-coding genes on the human X Chromosome, of which 221 of 823 (26.4%) are associated with an OMIM phenotype (p<0.05, hypergeometric test). Out of 5 human palindrome spacer genes, two are associated with OMIM phenotypes: *EMD* and *FLNA*, both broadly expressed genes found in the spacer of P24, which are associated with muscular dystrophy and neurological disorders, respectively (Table 1) (Bione et al. 1994, Fox et al. 1998, Clapham et al. 2012).

Despite their limited functional characterization, we find that palindrome gene families are well conserved across primates. All 17 gene families in human palindrome arms have at least one intact gene copy in chimpanzee, and 15 of 17 have at least one intact gene copy in rhesus macaque (Supplemental Table S5). Among spacer genes, 4 of 5 have at least one intact gene copy in chimpanzee, and 3 of 5 in rhesus macaque; the two genes with ascribed OMIM phenotypes are conserved in all three species (Supplemental Table S5). Three out of four arm and spacer gene families that are not conserved across all three species have paralogs with at least 85% protein identity elsewhere on the X Chromosome, which may reduce the impact of their loss. Gene families from palindromes shared by human, chimpanzee, and macaque also have conserved expression patterns: 20 of 21 such gene families have the same expression pattern in chimpanzee and human (Supplemental Fig. S7), and the same is true for 16 of 18 gene families conserved in rhesus macaque (Supplemental Fig. S8).

The conservation of palindrome gene families suggests that they are subject to purifying selection. Consistent with this, we found dN/dS values < 1.0 for 17 of 18 arm and spacer gene families conserved between all three species (Supplemental Table S6), twelve of which were significant using a likelihood ratio test (Supplemental Table S6). Nonetheless, the median dN/dS value for 18 X palindrome gene families is 0.36, compared to a median dN/dS value of 0.12 for protein-coding genes in the genome (Gayà-Vidal and Albà 2014, Biswas et al. 2016). Elevated dN/dS values could result from either relaxed purifying selection, or from positive selection at one or more sites. We therefore also performed a likelihood ratio test for positive selection across all 18 gene families, and found evidence of positive selection for 2 gene families (Supplemental Table S6). We note that with only three species for comparison, we were likely under-powered to detect positive selection, and our results should not be interpreted as evidence against positive selection in the other 16 gene families (Anisimova et al. 2001).

If palindrome gene families are subject to purifying selection, then we also predicted that they should be depleted for insertions and deletions (indels) between species. To define indels, we used a kmer-based method to identify stretches of at least 1 kb that lacked orthologous sequence in the other species (Fig. 5A). We then compared the fraction of bases falling within indels for protein-coding gene sequence (including exons, introns, and 1 kb upstream) versus other sequence. We performed this analysis individually for palindrome arms, palindrome spacers, and flanking sequence, revealing a significant depletion of indels within protein-coding gene sequence for all three regions (Fig. 5B). We conclude that natural selection has preserved protein-coding gene families in primate X-palindrome arms and spacers, despite their limited functional characterization in humans.

**Figure 5.**
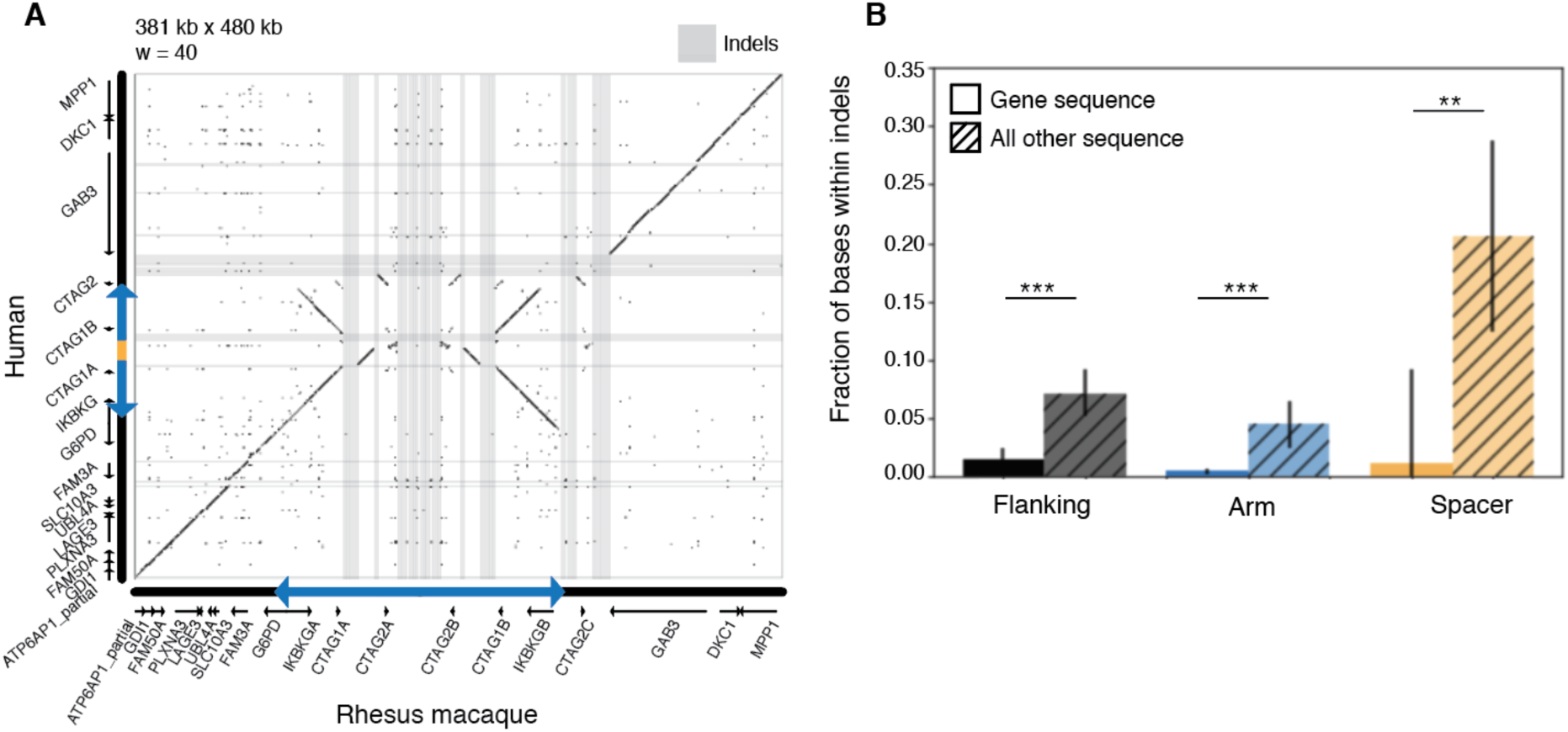
Natural selection has preserved X-palindrome gene families. Results based on 12 palindromes conserved between human, chimpanzee and rhesus macaque. a) Square dot plot comparing structure of P25 between human and rhesus macaque. Indels highlighted in gray; nearly all fall between protein-coding genes. b) Fraction of bases within indels for protein-coding gene sequence versus all other sequence. Results are the average of all pairwise species comparisons. Indels were defined as uninterrupted stretches of at least 1 kb in one species without orthologous sequence in the other species. **p<0.01, ***p<0.001, Mann-Whitney U.

### Enrichment of spacer deletions in human X-Chromosome palindromes

Having observed localized structural instability in X palindromes between primate species, we asked whether we could detect signatures of structural instability within the human population. To address this question, we used whole-genome sequencing data from the 1000 Genomes Project (The 1000 Genomes Project Consortium 2015). The 1000 Genomes Project dataset consists of short Illumina reads, which are not conducive to finding insertions or rearrangements. Instead, we asked whether we could detect X-palindrome spacer deletions, which we predicted would result in loss of sequence coverage over the palindrome spacer.

We searched for X-palindrome spacer deletions among 944 individuals from the 1000 Genomes Project. We limited our analysis to male samples because males have only one X Chromosome; this enabled us to analyze deletions among 944 X Chromosomes, and ensured that coverage depth over X-Chromosome deletions should be near zero. To identify palindrome spacer deletions, we screened for X Chromosomes with low normalized spacer coverage depth (Fig. 6A; also see Methods). We found four X Chromosomes with near-zero coverage in the spacer of P2 (Fig. 6A); visual inspection of coverage depth revealed that all four X Chromosomes had a deletion of about 25 kb, spanning not only the palindrome spacer but part of the inner palindrome arm (Fig. 6B). We performed the same analysis for the remaining 25 X palindromes, identifying a remarkable total of 149 palindrome spacer deletions across 9 different palindromes (Table 2; Supplemental Fig. S9).

**Figure 6.**
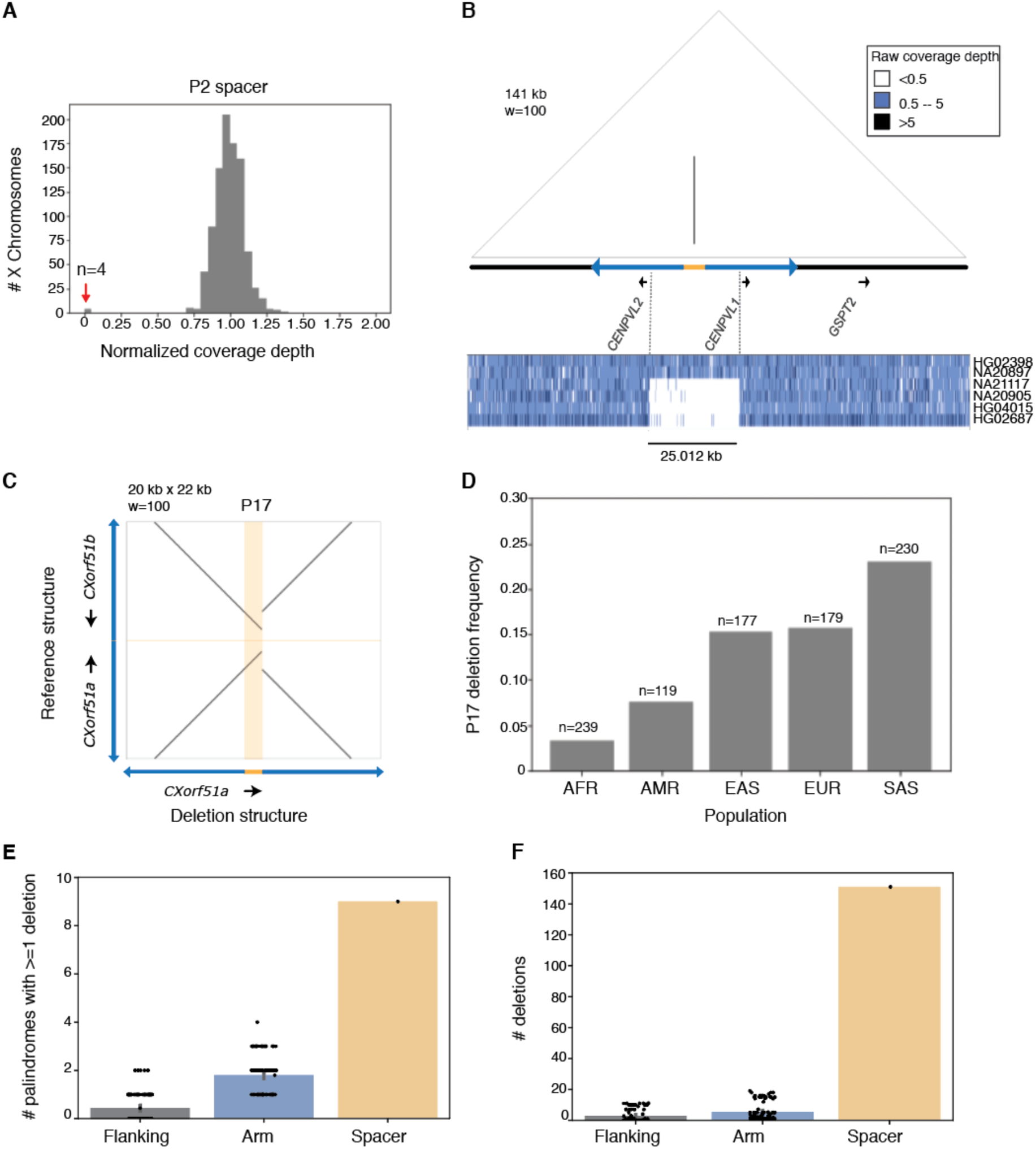
Deletions are enriched in human X-Chromosome spacers. a) Normalized coverage depths for P2 spacer. Red arrow indicates four X Chromosomes with depth near 0. b) Coverage depths across P2 and flanking sequence for two individuals with reference structure (HG02398, NA20897) and four with spacer deletions (NA21117, NA20905, HG04015, HG02687). c) Square dot plot comparing palindrome centers (spacer + 10 kb inner arm on each side) for P17 reference structure and P17 deletion. d) Frequency of P17 spacer deletions across five superpopulations from 1000 Genomes. EUR = European, AFR = African, AMR = Admixed Americas, EAS = East Asian, SAS = South Asian. e,f) Frequency of deletions detected in palindrome spacers compared to palindrome arms and flanking sequence. Size-matched regions from palindrome arms and single-copy sequence were selected at random; results from 100 iterations are shown.

**Table 2.** Palindrome spacer deletions among 944 X Chromosomes from the 1000 Genomes Project

| Palindrome | # X Chromosomes | Deletion size (kb) | Genes deleted | Breakpoint 1 | Breakpoint 2 |
| --- | --- | --- | --- | --- | --- |
| P17 | 126 | 3.36 | *CXorf51* (1/2 copies) | Arm | Arm |
| P8 | 14 | ~47 | *CXorf49* (1/2 copies) | Arm | Arm |
| P2 | 4 | 25.01 |  | Arm | Arm |
| P16 | 1 | ~587 | *LDOC1, SPANXC, SPANXA1* | Arm | Flanking |
| P11 | 1 | 9.93 |  | Spacer | Arm |
| P25 | 1 | 21.47 |  | Spacer | Arm |
| P5 | 1 | 17.89 |  | Arm | Arm |
| P22 | 1 | 8.46 | *PNMA6* (2/2 copies) | Arm | Arm |
| P1 | 1 | 3.97 |  | Arm | Arm |

For arm genes, the number of gene copies removed by the deletion is indicated in parentheses. All other genes are unique genes from the palindrome spacer or flanking sequence. Deletion sizes marked as approximate (~) were estimated from changes in coverage depth; all others are exact sizes based on split reads spanning the deletion breakpoint.

Although deletions were identified based on low depth of coverage in spacers, most breakpoints fell within palindrome arms (Fig. 6B, Table 2). In total, 145 of 944 X Chromosomes from the 1000 Genomes Dataset (15.4%) had a spacer deletion in at least one palindrome. Deletions ranged from 3 kb up to 587 kb in size, and in four cases removed one or more copies of a protein-coding gene family (Table 2). In two cases, breakpoints fell within tandem repeats, suggesting that they arose through NAHR (Supplemental Fig. S10). All other breakpoints fell within unique sequence, suggesting origins independent of NAHR (see Discussion). For the nine human X palindromes in which we identified at least one X Chromosome with a spacer deletion, we examined the structure of orthologous palindromes in chimpanzee. We found that sequence absent in one or more human X Chromosomes is present in the chimpanzee X Chromosome, confirming that the human structural polymorphisms result from deletions rather than insertions (Supplemental Fig. S11).

Out of a total of 149 palindrome spacer deletions, 126 were found in a single palindrome, P17 (Fig. 6C, Table 2). We wondered whether these represented independent deletion events, in which case we would expect different breakpoints in different X Chromosomes. Alternatively, if they represented a common structural polymorphism, all breakpoints should be identical. The deletion breakpoints for all 126 P17 deletions appeared similar by eye (Supplemental Fig. S12); we subsequently identified split reads from the 1000 Genomes Project spanning the same breakpoint for all 126 X Chromosomes, and further verified this shared breakpoint by PCR in five individuals selected at random (Supplemental Figs. S13, S14). We conclude that this deletion is a common polymorphism. This P17 deletion spans the palindrome’s spacer and inner arm, removing one copy of *CXorf51b*, a testis-expressed gene not associated with any phenotypes reported in OMIM (Table 1). The P17 spacer deletion is found in all five super-populations in 1000 Genomes, with frequencies ranging from 3% (Africa) to 23% (South Asia) of X Chromosomes (Fig. 6D). The 1000 Genomes dataset does not include phenotype information; however, we speculate that the rise of the P17 spacer deletion to high frequency is incompatible with strong reductions in viability or fertility, and that phenotypic effects, if any, are likely to be mild.

To determine whether genomic instability is elevated within palindrome spacers, we asked whether deletions were more common in palindrome spacers than in arms and flanking sequence. For each of 26 human X palindromes, we randomly selected regions of the arm and flanking sequence of the same size as the spacer, and we counted deletions by the criteria described above. For spacers, we observed deletions in 9 different palindromes; in contrast, we never observed more than 4 deletions in size-matched regions from palindrome arms and flanking sequence (p<0.01, bootstrapping analysis) (Fig. 6E). The difference was more dramatic in absolute terms: 149 deletions in spacers, with no more than 19 deletions in arms and flanking sequence (p<0.01, bootstrapping analysis) (Fig. 6F). We conclude that structural instability of X-palindrome spacers has persisted in our own species, and that one manifestation of this instability is deletions. Importantly, insertions and rearrangements are possible as well, but would not have been detected by our analysis.

### Two polymorphic human X-palindrome spacer deletions are not associated with azoospermia

Given that primate X-palindrome gene families are preserved by natural selection, we wondered whether deletions that remove one or more copies of human X-palindrome gene families negatively impact fitness. The two most common human X-palindrome spacer deletions from the 1000 Genomes Project each remove one copy of an uncharacterized testis-expressed gene family: *CXorf51* and *CXorf49*, respectively (Table 2). Since deletion of testis-expressed Y-palindrome gene families causes azoospermia (Vogt et al. 1996, Kuroda-Kawaguchi et al. 2001), we asked whether deletions of *CXorf51* and *CXorf49* are also associated with azoospermia, using a publicly available dataset containing capture-based targeted sequencing for 301 azoospermia cases and 300 normospermic controls (dbGaP ID: phs001023).

Selecting samples that met a minimum coverage threshold, we found that *CXorf51* deletions were equally prevalent in cases and controls: 47 deletions in 286 cases (16.4%), and 54 deletions in 292 controls (18.5%) (ns, Fisher exact test) (Supplemental Table S7). Interestingly, we also detected two deletions – both in controls – that appear to remove both copies of the *CXorf51* gene family. We found only one case with a *CXorf49* deletion and therefore cannot infer an association with azoospermia. To test for milder effects on spermatogenesis, we used PCR screening to identify *CXorf51* and *CXorf49* deletions in a set of 562 oligozoospermic men (sperm counts 0.1–20 million per cubic cm). We found *CXorf51* deletions in 68 men (12.1%), which is not significantly different from the percentage of men from the 1000 Genomes Project (13.3%, Fisher exact test), suggesting that *CXorf51* deletions are not enriched in oligozoospermic men (Supplemental Table S7). We identified a single *CXorf49* deletion in oligozoospermic men, which is a significantly lower rate than we observed among men from the 1000 Genomes Project (0.18% versus 1.4%, p<0.05, Fisher exact test). While our analyses do not support an association between *CXorf51* or *CXorf49* deletions and azoospermia or oligozoospermia, we cannot rule out more subtle defects in spermatogenesis, with resultant selection for retention of both gene copies.

## DISCUSSION

Massive palindromes are hallmarks of mammalian sex chromosomes, yet until now, there were few examples of sex chromosome palindromes that are conserved between species. We provide evidence that 12 palindromes have been conserved across three primate X Chromosomes for at least 25 million years, using a targeted sequencing protocol, SHIMS 3.0, that combines ultralong nanopore reads with a clone-based approach. Comparative genomic analyses of conserved X palindromes shed new light on palindrome evolution, including evidence that natural selection preserves understudied protein-coding genes within X-palindrome arms and spacers. We also report a novel structural instability of X-Chromosome palindromes: Rearrangements between species are concentrated around the center of symmetry, with a high frequency of palindrome spacer deletions observed among individuals from the 1000 Genomes Project.

The deep conservation of primate X palindromes has few parallels in the literature. Out of eight palindromic amplicons on the human Y Chromosome, only two were reported to have orthologous palindromes in rhesus macaque (Hughes et al. 2012). The genes *FLNA* and *EMD*, which are found in the human (X Chromosome) P24 spacer, are flanked by inverted repeats in 15 additional mammalian species, leading to the suggestion that these inverted repeats arose over 100 million years ago (Caceres et al. 2007). However, the inverted repeat sequence is not orthologous between all species, leaving open the possibility that they arose through independent duplications (Caceres et al. 2007). While the chimpanzee Y chromosome contains nineteen massive palindromes, fewer than half of them have homology to human Y palindromes, and abundant rearrangements between the human and chimpanzee Y chromosomes make it difficult to reconstruct the evolution of putative orthologs (Hughes et al. 2010). In contrast, all 12 palindromes that we found to be shared by human, chimpanzee, and macaque have clear orthology between arms and between flanking sequences, unambiguously establishing a common origin. Notably, each of these palindromes is found in a single copy on the X chromosome, distinguishing them from previous reports of co-amplified gene families on the X and Y chromosomes of Drosophila (Ellison and Bachtrog 2019), mouse (Cocquet et al. 2009, Cocquet et al. 2012, Soh et al. 2014), and bull (Hughes et al 2020). Our data provides strong evidence that palindromes can be maintained over tens of millions of years of evolution, in some cases with minimal structural change (e.g., P6; Supplementary Fig. 4). We note that 25 million years represents a lower bound on the age of the X-Chromosome palindromes, and that future high-resolution sequencing of mammalian X Chromosomes may reveal that these palindromes are conserved in more distantly related species.

Primate X palindromes could be conserved because they are inherently stable structures, or because they are preserved by natural selection. Although these explanations are not mutually exclusive, our results strongly favor natural selection. First, we demonstrate that palindromes are not inherently stable structures, exhibiting localized structural instability around the center of palindrome symmetry. Rearrangements are enriched around palindrome spacers and inner arms in structural comparisons of palindromes conserved between species; indeed, 7 of 12 palindromes shared by human, chimpanzee, and rhesus macaque have no spacer orthology between human and macaque. X-Chromosome palindromes remained structurally unstable during human evolution, as shown by a significant enrichment of deletions in palindrome spacers compared to palindrome arms or flanking sequence. Second, we provide multiple lines of evidence that palindrome gene families are targets of selection. Large (>1 kb) insertions and deletions in palindrome arms and spacers during the last 25 million years of primate evolution were depleted around protein-coding genes, and molecular analyses demonstrate purifying selection on protein-coding genes in X-palindrome arms and spacers. Notably, all twelve X palindromes conserved between human, chimpanzee, and rhesus macaque have at least one protein-coding gene in their arms or spacer. We conclude that palindromes are not inherently structurally stable, but rather are preserved through natural selection, most likely acting to preserve the integrity of protein-coding gene families.

The discovery of human X-palindrome spacer deletions in individuals from the 1000 Genomes Project, combined with evidence that purifying selection preserves human X-palindrome gene families, raises the question of whether human X-palindrome spacer deletions are pathogenic. To date, we are aware of one published report describing a pathogenic X spacer deletion. Periventricular nodular heterotopia (PNH), a common neurological disorder, is caused by loss-of-function mutations in *FLNA*, a broadly expressed spacer gene in P24 that encodes the actin crosslinking protein Filamin A (Fox et al. 1998). Although PNH is most frequently caused by missense mutations in *FLNA*, one affected family was found to have a 39-kb deletion spanning the P24 spacer and inner arm, removing spacer genes *FLNA* and *EMD* (Clapham et al. 2012). Although we found spacer deletions in nine different human X palindromes across individuals from the 1000 Genomes Project, none of the deletions occurred in P24. Given that the 1000 Genomes Project does not include phenotype information, we do not know whether any of the spacer deletions we observed are pathogenic. However, 4 of 9 human X spacer deletions removed at least one copy of a protein-coding gene family, and two of these removed all functional copies of a gene family (*PNMA6*: 2/2 copies; *LDOC1:* 1/1 copies). We speculate that deletions that remove one or more X-palindrome genes may result in mildly deleterious phenotypes, such as subtle defects in spermatogenesis; this hypothesis is consistent with our observation that insertions and deletions affecting X-palindrome genes are depleted between species.

NAHR is a common cause of palindrome rearrangements on the human X and Y Chromosomes (Lakich et al. 1993, Small et al. 1997, Aradhya et al. 2001, Lange et al. 2009, Scott et al. 2010). However, out of the nine X-palindrome spacer deletions we observed in males from the 1000 Genomes Project, only two displayed breakpoint homology consistent with NAHR. One possible explanation for non-NAHR based rearrangements is replication errors: Duplication of a single-copy human X gene near P24 has been reported to result from Fork Stalling and Template Switching (FoSTeS), in which replication machinery repeatedly stalls within a low-copy repeat and switches templates, creating a rearrangement of the form duplication-inverted triplication-duplication (Carvalho et al. 2009, Carvalho et al. 2011). FoSTeS rearrangements lead to increased copy number, yet replication errors in different genomic contexts could lead to sequence loss. Indeed, intra-strand pairing and subsequent replication slippage were proposed to explain deletions within a 15-kb transgenic palindrome in mice, which underwent large asymmetric deletions around the center of palindrome symmetry within a single generation (Akgun et al. 1997). While replication errors represent one plausible explanation for primate X-palindrome spacer deletions, future studies will be required to rule out other mechanisms.

Our work affirms the importance of high-quality genome assemblies for comparative genomics, by revealing a wealth of conserved X palindromes with signatures of natural selection that were largely missing from existing chimpanzee and rhesus macaque X-Chromosome assemblies. In contrast to other genome assembly methods, SHIMS 3.0 enables the verification of palindromes and other genomic structures through the generation of multiple full-length nanopore reads from the same clone. In recent years, long-read technologies have also been incorporated into mammalian genome assemblies generated using a whole-genome shotgun assembly approach (see Gordon et al. 2016, Bickhart et al. 2017, Low et al. 2019, Miga et al. 2020, and others). While long-read WGS assemblies offer substantial improvements over short-read WGS assemblies, we nevertheless found that the fraction of primate X palindromes represented accurately in two long-read WGS assemblies hovered around 50%, demonstrating that clone-based methods remain necessary to confidently resolve complex genomic structures. Indeed, a recent nanopore WGS assembly of the human genome achieved greater continuity than the previous human genome assembly, yet was still missing nearly 20% of segmental duplications and other hard-to-sequence regions of the genome that had been previously sequenced using large-insert clones (Miga et al. 2020).

Going forward, we propose that long-read whole-genome shotgun assemblies and SHIMS 3.0 may be used in tandem to improve the representation of palindromes in mammalian genomes. Our study used synteny between primate X Chromosomes to identify candidate regions in chimpanzee and rhesus macaque that were likely to contain palindromes; candidate regions were then targeted with SHIMS 3.0 to generate finished sequence. We propose that long-read WGS assemblies may serve a similar role: In our own comparison of primate X-Chromosome assemblies, we found that while some palindromes were missing entirely from long-read WGS assemblies, the majority were present but incomplete. Long-read WGS assemblies may thus serve as a guide for identifying the positions of putative palindromes, which can then be finished using SHIMS 3.0 or other clone-based approaches. Future comparative analyses using high-resolution sequence will reveal whether conserved palindromes are a feature of other mammalian X Chromosomes, and if so, shed further light on the balance of structural instability and natural selection that govern their evolution. High-resolution X chromosome sequence for other great apes (gorilla and orangutan) may be particularly useful for probing the dynamics of X palindrome spacer inversions and deletions.

The conserved X-Chromosome palindromes that we describe here represent a substantially understudied class of genomic sequence. The technical challenges presented by palindromes go beyond the generation of accurate reference sequence: Many common bioinformatics tools, such as those used for quantifying gene expression or detecting mutations, automatically discard multi-mapping reads, rendering palindromes invisible in downstream analyses (Godfrey et al. 2020). The issue can be overcome by selection of tools like kallisto that probabilistically assign multi-mapping reads (Bray et al. 2016), yet this is not routinely done for large-scale genomic analyses. Given these challenges, it is not surprising that human X-palindrome gene families remain poorly characterized compared to other human X-linked genes (see also Mueller et al. 2013). Many human X-palindrome genes are classified as cancer-testis antigens (CTAs), genes defined by expression in the testis as well as in cancerous tumors, yet the mechanisms and significance of this phenomenon are not well understood (Simpson et al. 2005). Deletion or inversion of a single arm of a palindrome containing a testis-specific gene family in mouse yielded no observable phenotypes, leading to the suggestion that palindrome structures may primarily have benefits over longer evolutionary timescales, perhaps through purging deleterious mutations or fixing beneficial mutations through rapid gene conversion (Kruger et al. 2018). We propose that palindromes have a fundamentally different biology than unique sequence – a biology that does not readily align with our expectations or our standard methods of imputing function, including murine mouse models or association with Mendelian disease, both of which depend on observing a strong phenotype within a single generation. Technical advances that facilitate the study of X palindromes and other amplicons will be essential to illuminate their biology, with implications for X-Chromosome evolution as well as human health and disease.

## METHODS

### Palindrome annotation

Candidate regions likely to contain human X-Chromosome palindromes were identified from the genomicSuperDups track in the UCSC Genome Browser (Kent et al. 2002) using the following criteria: Inverted repeats >8 kb in length and displaying >95% sequence identity, with <500 kb between arms. For each candidate region, we divided the sequence into overlapping 100-base-pair windows, then aligned these windows back to the candidate region using bowtie2, with settings to return up to 10 alignments with alignment scores >-11 (Teitz et al. 2018). We then created a bedGraph file for each candidate region in which the value for each position represents the number of times the window starting at that position aligns to that region, and visualized the bedGraph file using the Interactive Genome Viewer (IGV) (Robinson et al. 2011). Putative palindromes boundaries were annotated manually based on the start and end of long stretches of multi-mapping windows, and filtered for arms >8 kb and arm-to-arm identity >99%. The same method was used to annotate palindromes within chimpanzee and rhesus macaque SHIMS 3.0 assemblies for regions orthologous to human X palindromes.

### Sequence alignments and dot plots

Square and triangular dot plots were generated using custom Perl code (http://pagelabsupplement.wi.mit.edu/fast_dot_plot.pl). Unless otherwise noted, sequence alignments were performed using ClustalW with default parameters (Thompson et al. 1994). To identify and exclude regions of poor alignment, ClustalW sequence alignments were scanned using a sliding 100-bp window and filtered to exclude windows with fewer than 60 matches between species, using custom Python code.

### Human gene annotation

GENCODE 34 gene annotations for the human X Chromosome were downloaded from ENSEMBL using the BioMart package in R. Annotations were filtered for protein-coding genes only, and the APPRIS principal transcript was selected for each gene. If there were multiple principal transcripts, the longest principal transcript was selected, and if there were multiple principal transcripts of equal length, the longest principal transcript with the highest transcript support level (TSL) was selected. There were three exceptions as follows:

1. There were two protein-coding genes with the same ENSEMBL gene name: ENSG00000158427 and ENSG00000269226, both named *TMSB15B*. We refer to them as *TMSB15BA* and *TMSB15BB*, respectively.
2. The palindrome arm genes *PNMA6B* and *AC152010.1* were included despite being annotated as pseudogenes, because each gene was annotated as having a protein-coding paralog in the other arm (see Supplemental Note S1).
3. The principal transcript for palindrome arm gene *TCP11X2* encoded a different protein than the principal transcript for its paralog *TCP11X1*. For consistency, the isoform encoding the longer protein was selected as the principal transcript for both; transcript ENST00000642911 (marked “alternative2”) was therefore used for *TCP11X2*.

### Human gene expression

Gene expression was calculated for a subset of samples from the GTEx project (v8) as follows. For each tissue subtype in GTEx, 5-10 of the highest quality samples were selected based on a combination of RNA integrity (RIN), mapped-read library size, and intronic read mapping rate. BAM files containing all reads (mapped and unmapped) from these samples were accessed through Terra (https://app.terra.bio), and used to generate FASTQ files. Transcript expression levels in TPM were estimated using kallisto with sequence-bias correction (--bias) using GENCODE 34 gene annotations, then summed to obtain gene expression levels. Results were filtered to include protein-coding genes only and TPM values were re-normalized to 1 million for each sample. For human X-palindrome arm gene families, expression levels for both arm genes were averaged to return gene family expression levels. To analyze expression of human X palindrome gene families in spermatogenesis, we downloaded publicly available SRA files from Jan et al. 2017 (SRP069329) and analyzed them with kallisto as described above.

### Transposable elements

We analyzed transposable element density using RepeatMasker (https://www.repeatmasker.org) with default settings.

### Clone selection and sequencing

All chimpanzee clones selected for sequencing were from BAC library CH251 (https://bacpacresources.org), which derives from a single male individual (“Clint”) used in initial sequencing of the chimpanzee genome (Chimpanzee Sequencing and Analysis Consortium 2005). All rhesus macaque clones selected for sequencing were from BAC library CH250, which derives from a single male individual of Indian origin (https://bacpacresources.org). Sequencing was performed using the SHIMS 3.0 protocol (Bellott et al 2020). Regions covered by one or more nanopore reads, but no Illumina reads, were marked as “problem regions” and excluded from downstream analysis; these regions represented <1% of all sequence generated for this project. Our assemblies also include sequence from 7 chimpanzee clones previously sequenced and deposited in GenBank; these clones each contained part or all of a palindrome arm, but did not contain internal repeats, making them suitable for assembly without long reads (Supplemental Table S8).

### Comparison to existing X-Chromosome assemblies

X-Chromosome assemblies for Pan_tro_3.0 (CM000336.3), Mmul_8.0.1 (CM002997.3), and Mmul_10 (CM014356.1) were downloaded from Ensembl (www.ensembl.org). The X-Chromosome assembly for Clint_PTRv2 (CM009261.2) was downloaded from NCBI (https://www.ncbi.nlm.nih.gov). Regions orthologous to each chimpanzee and rhesus macaque palindrome, as identified in SHIMS 3.0 assemblies, were extracted from each X-Chromosome reference assembly using custom Python code. We generated triangular dot plots for each extracted region, and square dot plots comparing each extracted region to the orthologous SHIMS 3.0 assembly. Categorizations were made as follows: 1) “Missing” = no palindrome, 2) “Incomplete” = a palindrome was partially present but misassembled, and 3) “Accurate” = palindrome arms and spacer aligned fully to the orthologous SHIMS 3.0 assembly.

### Primate gene annotation

Primate gene annotations were performed manually using alignment of human exons and alignment of testis RNA-Seq reads for guidance. For each SHIMS 3.0 assembly from chimpanzee and rhesus macaque, we aligned exons from the corresponding human genomic region using BLAT (Kent 2002), and verified that splice sites were conserved (acceptor: AG; donor: GT or GC). In instances where splice sites were not conserved, we aligned testis RNA-Seq from the appropriate species. If we identified reads supporting the existence of an alternative splice site, we selected the alternative splice site; otherwise, we selected the original position and annotated the transcript as a pseudogene. In a subset of cases, part or all of the transcript fell into a problem area supported by nanopore reads but not Illumina reads. In these instances, gene annotations were modified using alignment of testis RNA-Seq and/or whole-genome sequencing (WGS) Illumina reads from a single male of the appropriate species. Chimpanzee testis RNA-Seq: SRR2040591, Rhesus macaque testis RNA-Seq: SRR2040595. Chimpanzee WGS Illumina: SRR490084 and SRR490117; Rhesus macaque WGS Illumina: SRR10693566.

### Primate gene expression

The latest transcriptomes for chimpanzee and rhesus macaque were downloaded from ensembl.org (Pan_troglodytes.Pan_tro_3.0.cdna.all.fa and Macaca_mulatta.Mmul_10.cdna.all.fa, respectively), and merged with newly annotated palindrome arm and spacer genes from SHIMS 3.0 assemblies. To prevent redundancy between our transcripts and transcripts representing the same genes that were already present in existing transcriptomes, we used BLAST to identify and remove existing transcripts that aligned to newly annotated genes over >50% of their length and with >95% sequence ID. Gene expression was calculated using RNA-Seq reads from the following publicly available datasets containing at least 5 different tissues including testis: Chimpanzee, Brawand et al, 2011; Rhesus macaque, Merkin et al. 2012. Transcript expression levels were calculated using kallisto with sequence-bias correction (--bias), and summed to gene expression levels. To enable comparison of expression between conserved human and primate gene families, all primate X-palindrome genes were grouped based on their closest human X-palindrome gene family, and gene family expression levels were calculated accordingly.

### Definition of orthologous palindromes

For each palindrome identified in chimpanzee or rhesus macaque SHIMS 3.0 assemblies, we generated alignments between Arm 1 of the non-human primate and Arm 1 of the putative human ortholog. Orthologous palindromes were required to meet two criteria, designed to establish an unambiguous common origin. First, at least 20% of the non-human primate palindrome arm was required to align to the putative human ortholog. Second, the alignable portion of the human palindrome arm was BLASTed against the complete non-human primate region (including palindrome arms, spacer, and flanking sequence) using default parameters. More than 90% of positions in high-quality BLAST hits (>1 kb, 95% for chimpanzee versus human; >1 kb, 90% for rhesus macaque versus human) were required to map to the palindrome arms.

### Calculation of divergence

Divergence was calculated by generating pairwise alignments using ClustalW, then calculating p-distance with MEGA X (Kumar et al. 2018). For alignment of arms between species, we generated pairwise alignments using Arm 1 from each species.

### Calculation of fraction orthologous sequence

Pairwise alignments between species were generated as described above. The fraction of orthologous sequence was calculated as (total bases in unfiltered alignment windows)/(total bases in starting sequence), after excluding bases from problem areas.

### Analysis of Online Mendelian Inheritance in Man (OMIM) phenotypes

We downloaded the genemap2.txt file from the OMIM database (https://www.omim.org/) (McKusick-Nathans Institute of Genetic Medicine 2020). We filtered for phenotypes linked to a single X-linked gene using custom Python code, and calculated the fraction of all protein-coding X-linked genes with an OMIM phenotype, relative to the fraction of X-palindrome arm genes and X-palindrome spacer genes with an OMIM phenotype.

### Calculation of dN/dS

Alignments of coding sequence from X-palindrome arm and spacer genes conserved between human, chimpanzee, and rhesus macaque were performed using default parameters from ClustalW. dN/dS values were calculated using the basic model in PAML (model = 0, NSsites = 0) (Yang 2007). To test the significance of calculated dN/dS values, we compared the likelihood of calculated values against a model where dN/dS was fixed at one. To test for positive selection, we compared the likelihood of model M1a (neutral evolution) versus model M2a (positive selection at one or more sites). We defined significance for both comparisons using the chi-squared distribution and appropriate degrees of freedom.

### Depletion of indels within protein-coding genes

Sequence from one species was broken into overlapping kmers with step size=1 and aligned to orthologous sequence from the other species using bowtie2 with settings to return up to 10 alignments with alignment scores >-11. Kmer size was either 100 (human-chimpanzee comparisons) or 40 (human-rhesus macaque comparisons). Indels were defined as stretches of at least 1 kb from one species that had no aligned kmers from the other species.

### 1000 Genomes data analysis

We analyzed whole-genome sequencing data from 1225 males from the 1000 Genomes Project (1000 Genomes Project Consortium 2015). Data selection, sequence alignment, GC bias correction, and repeat masking were performed as described in Teitz et al. 2018. We calculated average read depth for palindrome arms, spacers, and flanking sequence, normalized to a 1-Mb region of the X Chromosome without palindromes, using custom Python scripts. We filtered for males whose X Chromosome normalization region had an average read depth >=2, restricting our downstream analysis to 944 males. For small spacers (<3 kb), we expanded the area over which we calculated average read depth symmetrically into the inner palindrome arm until reaching 3 kb.

To identify candidate spacer deletions, we initially filtered for palindrome spacers with a normalized read depth below 0.25. After visualizing histograms of spacer depth across 944 males for each palindrome, we noticed a second peak centered around 0.25 for P17. To include all candidate P17 spacer deletions, we therefore raised our initial filtering threshold for P17 to 0.5. Read depths for all candidate spacer deletions were viewed using IGV. Candidate deletions that did not have a clear reduction in read depth in the spacer were excluded; all others were included in Table 2.

### Identification of split reads spanning deletion breakpoints

Forward and reverse reads from individuals with deletions were aligned separately to the region of the suspected deletion with settings to return up to 10 matches of minimum alignment score −11. We identified read pairs in which one read aligned and the other did not, and returned the sequence of the read that did not align. In the case of P17 deletions, we inspected unaligned reads by eye from four males to identify reads spanning the breakpoint. Finding that all of these males had the same breakpoint, we then used this breakpoint (+/- 10 base pairs on each side) to screen unmapped reads from all other males with suspected P17 deletions. For individuals where a breakpoint read could not be found using the primary 1000 Genomes dataset reads (1000genomes.sequence.index), we used a deeper 1000 Genomes dataset (1000G_2504_high_coverage.sequence.index).

### PCR verification of human palindrome spacer deletions

Patient genomic DNAs were purchased from Coriell Cell Repositories (HG01872, HG02070, HG02398, HG02687, HG03295, HG04015, HG04219, NA11919, NA18645, NA19086, NA19652, NA20351, NA20897, NA20905, NA21116, NA21117, NA21133). DNAs were tested for the presence or absence of palindrome spacers using primer pairs described in Supplemental Table S9. PCR was performed using 50 ng of DNA as template in a total volume of 20 μl (10 mM Tris– HCl [pH 9], 1.5 mM MgCl2, 50 mM KCl, 0.1% Triton X-100, 0.2 mM dNTPS, 0.5 μM primers, 0.5 U Taq polymerase). PCR cycling conditions for all primers were as follows: 94°C (30 s), 61°C (30 s), 72°C (1 min) for 35 cycles. Long range PCR was performed using Advantage 2 Polymerase following the manufacturer’s protocol (Clontech Laboratories, Mountain View CA).

### Association of *CXorf51* and *CXorf49* deletions with azoospermia

Palindrome spacer deletions removing one copy of *CXorf51* or *CXorf49* in dbGAP dataset phs001023 were detected based on reduced coverage depth, as described above for 1000 Genomes, with modifications as follows. Dataset phs001023 was generated using target-based capture sequencing rather than whole-genome shotgun sequencing, making it inappropriate for *de novo* deletion discovery. We therefore selected coordinates for detection of *CXorf51* and *CXorf49* spacer deletions based on three criteria: 1) coordinates overlap part or all of the deletion identified from the 1000 Genomes analysis, 2) coordinates contain at least 2 targeted probes, and 3) coverage depth within coordinates is predicted to decrease by >50% when the deletion identified from 1000 Genomes is present.

### Association of *CXorf51* and *CXorf49* deletions with oligozoospermia

We analyzed 562 DNA samples from oligozoospermic men previously collected by our lab. We excluded samples from men with Y chromosome deletions, varicocele, undescended testicles, or other known risk factors for oligozoospermia. Palindrome spacer deletions removing one copy of *CXorf51* or *CXorf49* were detected using the same primers and PCR conditions used for verification of deletions from 1000 Genomes (see above). Each DNA sample was tested using one set of primers expected to yield no product in samples with the deletion (P17 inner arm, P8 inner arm) and one set of primers expected to yield a specific product in samples with the deletion (P17 breakpoint, P8 breakpoint) (Supplemental Table S9).

### Human data

These studies were approved by the Massachusetts Institute of Technology’s Committee on the Use of Humans as Experimental Subjects. Informed consent was obtained from all subjects.

## DATA ACCESS

BAC sequences generated in this study have been submitted to GenBank (https://www.ncbi.nlm.nih.gov/) under accession numbers AC280414 through AC280580 (Supplemental Table S10). Codes for replicating these analyses are included in Supplemental Code as well as on GitHub (https://github.com/ejackson054/primate-X-palindromes).

## ACKNOWLEDGEMENTS

We thank A.K. San Roman and J.L. Mueller for comments on the manuscript, and A.K. Godfrey and other current and former Page lab members for valuable advice throughout the project. We thank S. Silber, R. Oates, M.C. Summers, V. Cardone, P. Patrizio, B. Gilbert, W.A. Hogge, C. Brunning, M. Jamehdor, B. Shapiro, K. Monaghan, J.C. Petrozza, L. Oman-Ganes, and J. Roberson for patient samples. This work was supported by the Howard Hughes Medical Institute, the Whitehead Institute, and generous gifts from Brit and Alexander d’Arbeloff and Arthur W. and Carol Tobin Brill.

## Author contributions

E.K.J., D.W.B, H.S., J.F.H., and D.C.P designed the study. T.-J.C. generated nanopore and Illumina sequencing reads. E.K.J. and D.W.B. assembled and finished clones. T.P. performed PCR experiments. E.K.J. performed computational analyses with assistance from D.W.B. and H.S. E.K.J. and D.C.P. wrote the manuscript.

