## Supplemental Materials for "Large palindromes on the primate X Chromosome are preserved by natural selection"

**Table of Contents**

**Supplemental Note 1:** Treatment of human X-palindrome genes with conflicting annotations

**Supplemental Figure 1:** Expression of human X-palindrome gene families
**Supplemental Figure 2:** Expression of human testis-biased X-palindrome gene families during spermatogenesis
**Supplemental Figure 3:** Structural comparisons between palindromes in SHIMS 3.0 assemblies and existing X-Chromosome assemblies
**Supplemental Figure 4:** Definition of orthologous palindromes
**Supplemental Figure 5:** Annotated square and triangular dot plots of primate X palindromes
**Supplemental Figure 6:** Additional examples of spacer configurations in orthologous palindromes
**Supplemental Figure 7:** Expression of gene families from palindromes shared by human, chimpanzee, and macaque in chimpanzee
**Supplemental Figure 8:** Expression of gene families from palindromes shared by human, chimpanzee, and macaque in macaque
**Supplemental Figure 9:** Normalized coverage depths for eight palindrome spacers with at least one deletion in the 1000 Genomes dataset
**Supplemental Figure 10:** Human spacer deletions with breakpoints within tandem repeats
**Supplemental Figure 11:** Structural comparisons between human reference, human deletion, and chimpanzee for nine X palindromes with spacer deletions
**Supplemental Figure 12:** Coverage depth for males with P17 spacer deletions
**Supplemental Figure 13:** Junction for P17 spacer deletion
**Supplemental Figure 14:** Verification of human X-palindrome spacer deletions

**Supplemental Table 1:** Coordinates of human X palindromes in hg38
**Supplemental Table 2:** Clones sequenced for this project
**Supplemental Table 3:** Status of palindromes identified in SHIMS 3.0 in other chimpanzee X assemblies
**Supplemental Table 4:** Status of palindromes identified in SHIMS 3.0 in other rhesus macaque X assemblies
**Supplemental Table 5:** Conservation of X-palindrome gene families
**Supplemental Table 6:** Purifying selection on X-palindrome genes
**Supplemental Table 7:** Chimpanzee clones used in this project that were previously sequenced and deposited in GenBank
**Supplemental Table 8:** Rates of P17 spacer deletions in azoospermic and oligozoospermic men
**Supplemental Table 9:** PCR primers for gels shown in Supplemental Figure 14
**Supplemental Table 10:** Chimpanzee and rhesus macaque clones sequenced for this project and deposited in GenBank

**Supplemental Note 1:** Treatment of human X-palindrome genes with conflicting annotations

There were three instances in which a gene in one arm of a palindrome was designated as protein-coding while the homologous sequence in the other arm was designated a pseudogene: *IKBKG* (protein-coding) and *IKBKGP1* (unprocessed pseudogene); *PNMA6A* (protein-coding) and *PNMA6B* (unprocessed pseudogene), and *AC236972.4* (protein-coding) and *AC152010.1* (processed pseudogene). We decided whether to include gene copies marked as pseudogenes in downstream analyses, i.e., whether their expression should be averaged with that of the corresponding protein-coding gene, as follows:

1) *PNMA6A* encodes a protein of 399 amino acids. *PNMA6A* and *PNMA6B* differ in their coding sequence by only a single missense substitution. The 3' UTR of *PNMA6B* is truncated, but the significance of this is unclear. Given that *PNMA6B* encodes an intact protein-coding sequence, we chose to include *PNMA6B* in downstream analyses.

2) *AC236972.4* encodes a protein of 2061 amino acids. *AC152010.1* has a nonsense substitution, but contains a downstream start codon that would lead to translation of the terminal 1253 amino acids of *AC236972.4.* Given that *AC152010.1* encodes a protein encompassing more than half the length of the original protein, we chose to include *AC152010.1* in downstream analyses.

3) *IKBKG* encodes a protein of 419 amino acids. *IKBKGP1* is a well-characterized pseudogene lacking the promoter and first four exons of *IKBKG* (Aradhya et al. 2001); we therefore chose not to include it in downstream analyses.

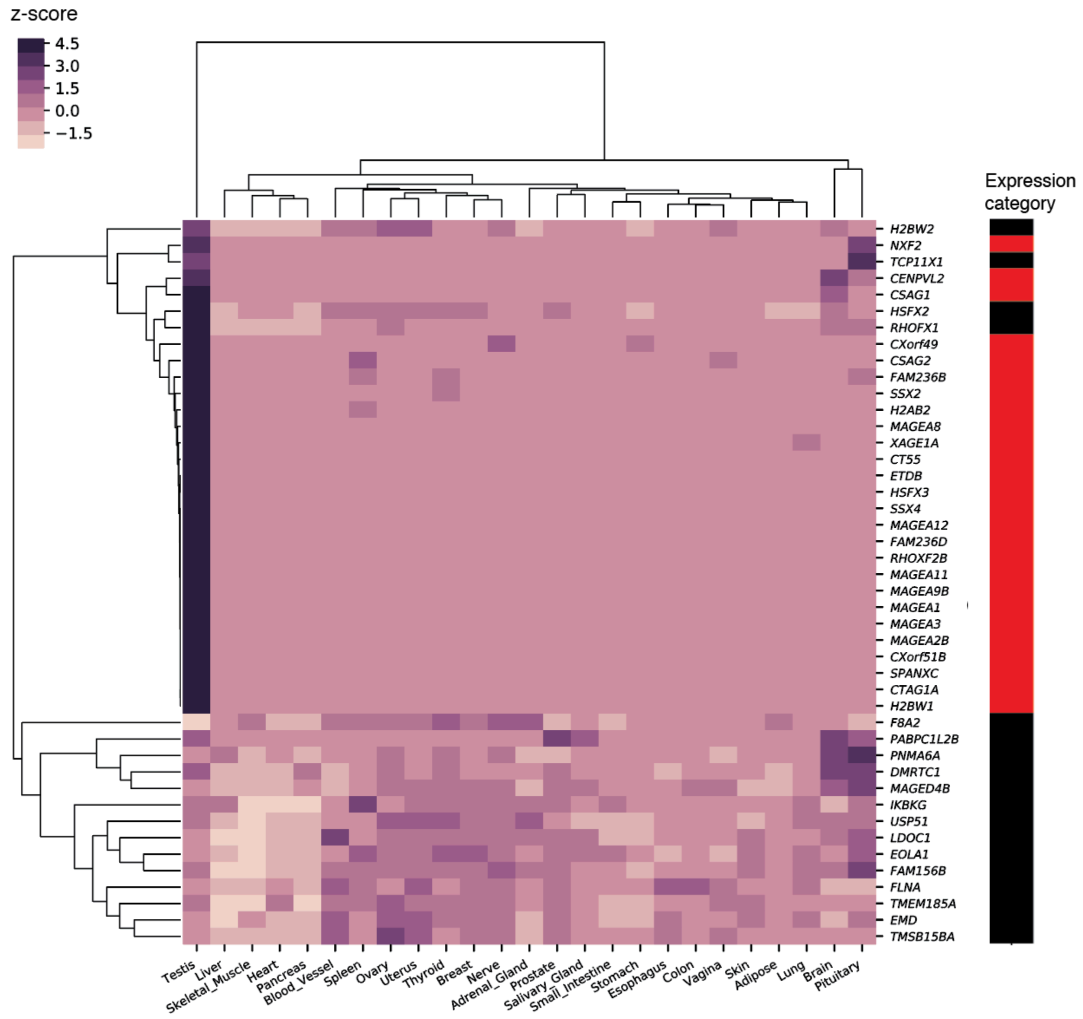


**Supplemental Figure S1.** Expression of human X-palindrome gene families. Heatmap shows z-score for each expressed gene (>2 TPM in at least one tissue) across 25 tissues from GTEx, with row and column order determined by hierarchical clustering. Expression category: Shows whether expression is testis-biased (red) or broad (black). Testis-biased: Minimum 2 TPM in testis, and testis accounts for >25% of log2 normalized expression summed across all tissues. Broad: All other expressed genes.

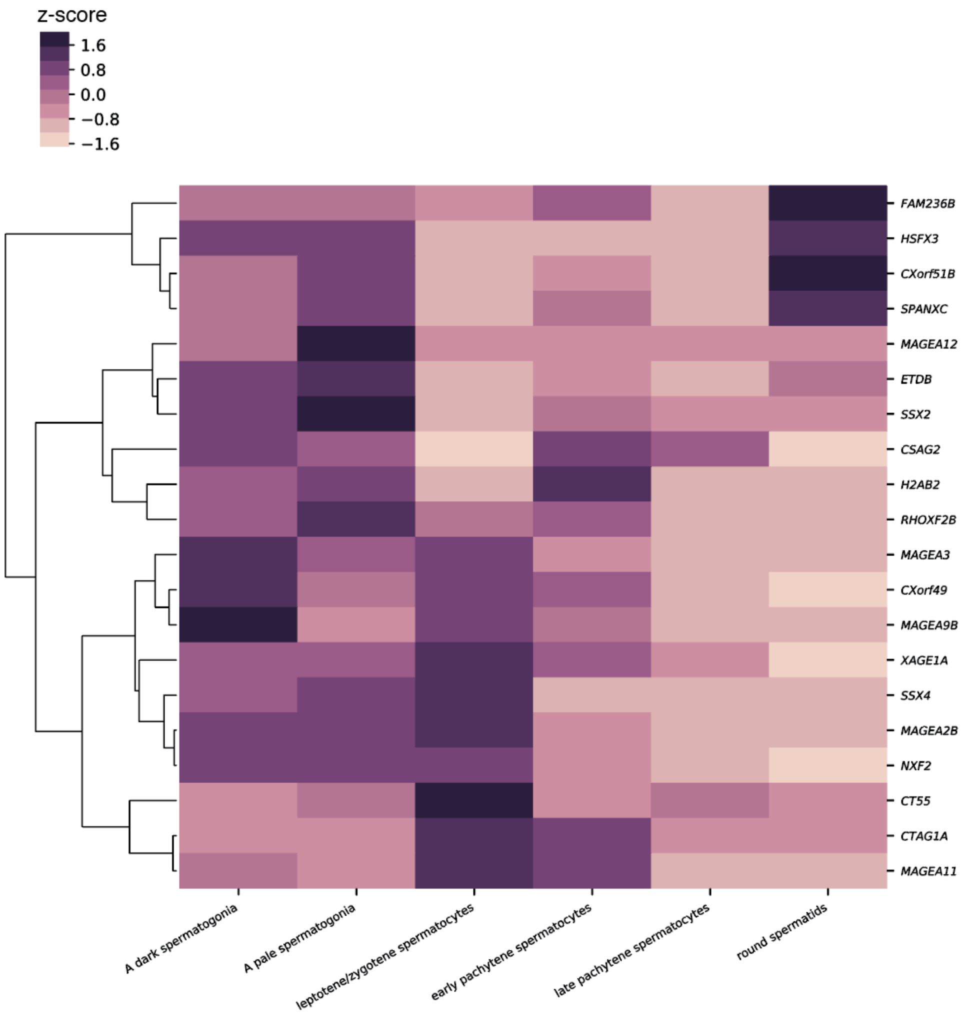


**Supplemental Figure S2.** Expression of human testis-biased X-palindrome gene families during spermatogenesis. Heatmap shows z-score for each expressed gene (>2 TPM in at least one spermatogenic stage) across six spermatogenic stages from Jan et al. 2017, with row order determined by hierarchical clustering.

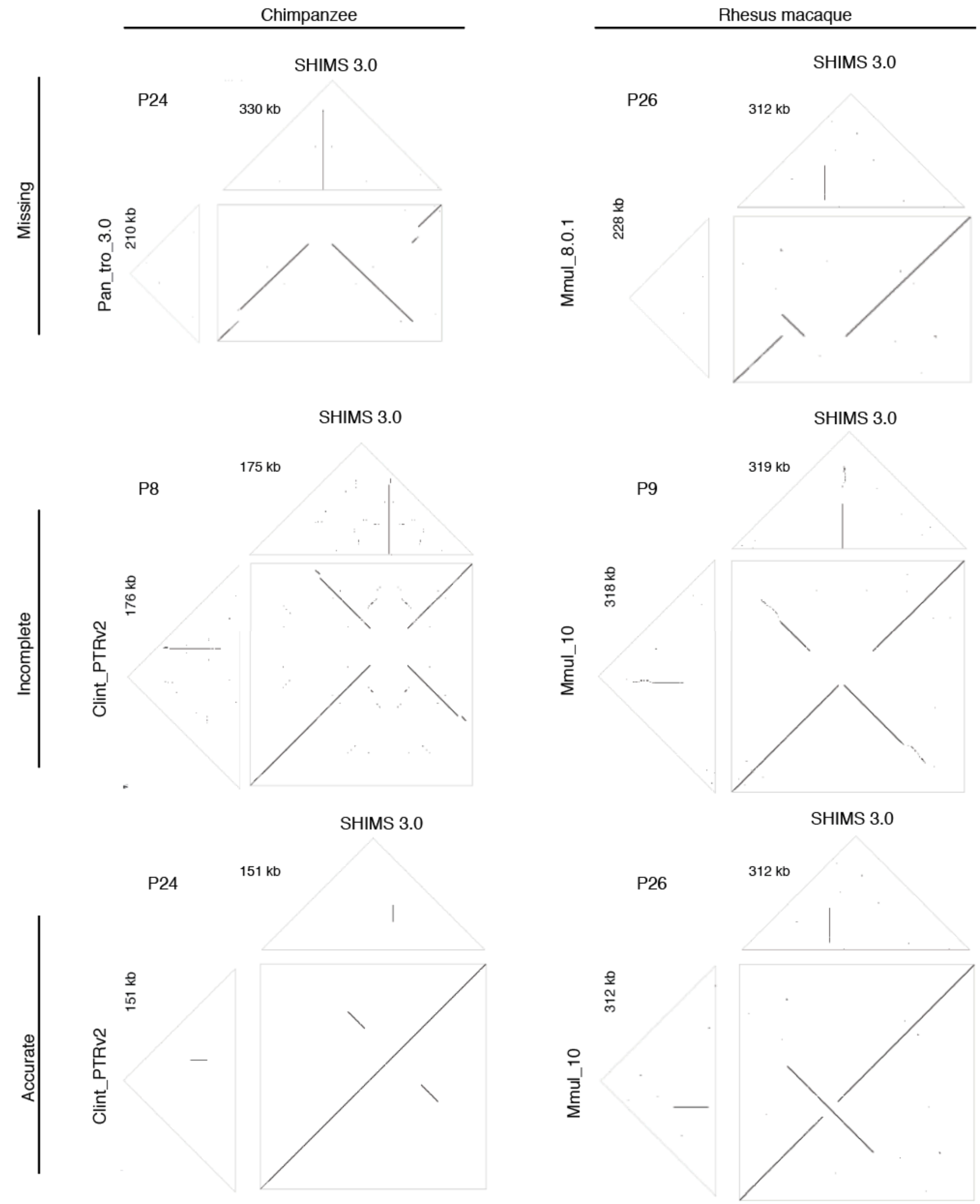


**Supplemental Figure S3.** Structural comparisons between palindromes in SHIMS 3.0 assemblies and existing X-Chromosome assemblies. Missing: No palindrome present in non-SHIMS 3.0 assembly. Incomplete: Part of palindrome present in non-SHIMS 3.0 assembly. Accurate: Full palindrome present in non-SHIMS 3.0 assembly. For accurate palindrome assemblies, note the presence in the square dot plot of an uninterrupted diagonal line and two arms that each map twice to the SHIMS 3.0 assembly. w=100 for all triangular and square dot plots.

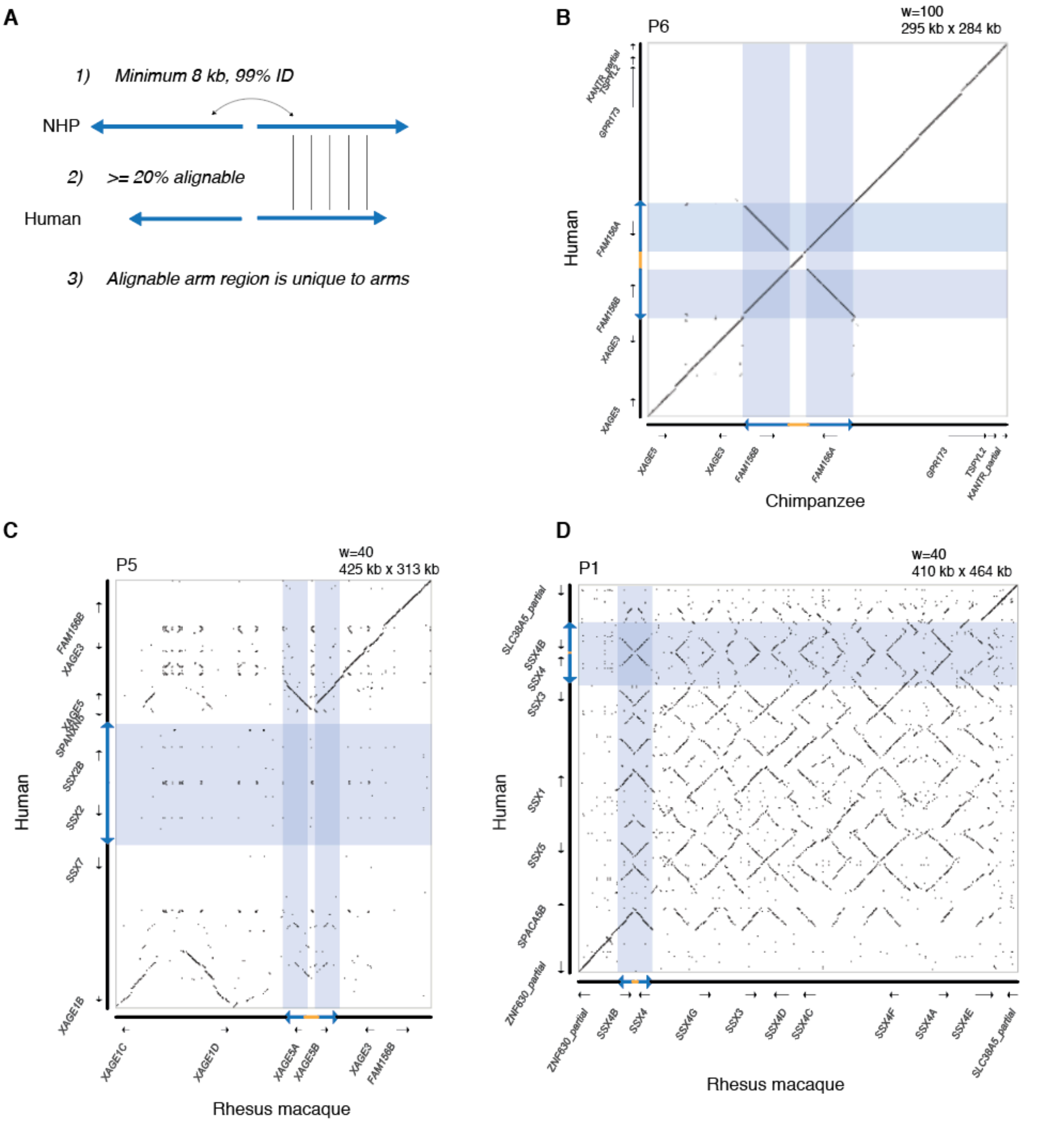


**Supplemental Figure S4.** Definition of orthologous palindromes. a) Criteria for defining orthologous palindromes. NHP = non-human primate (chimpanzee or rhesus macaque). Human and NHP palindrome arms were aligned with ClustalW, and required to have at least 20% alignment between species. Palindromes were excluded if the alignable region between palindrome arms mapped equally well to flanking sequence using reciprocal BLAST hits (>10% positions in high-quality hits mapping outside of palindrome arms). b) Example of an orthologous palindrome. Human palindrome arms map exactly twice to chimpanzee palindrome arms, and vice versa. c, d) Examples of palindromes that are not orthologous. c) Human palindrome arms have no orthologous sequence in rhesus macaque. Rhesus macaque palindrome arms correspond to flanking sequence in human. d) Rhesus macaque palindrome arms correspond equally well to more than two positions in human, and vice versa. Note that the region with the strongest orthology to rhesus macaque palindrome arms corresponds to flanking sequence in human.

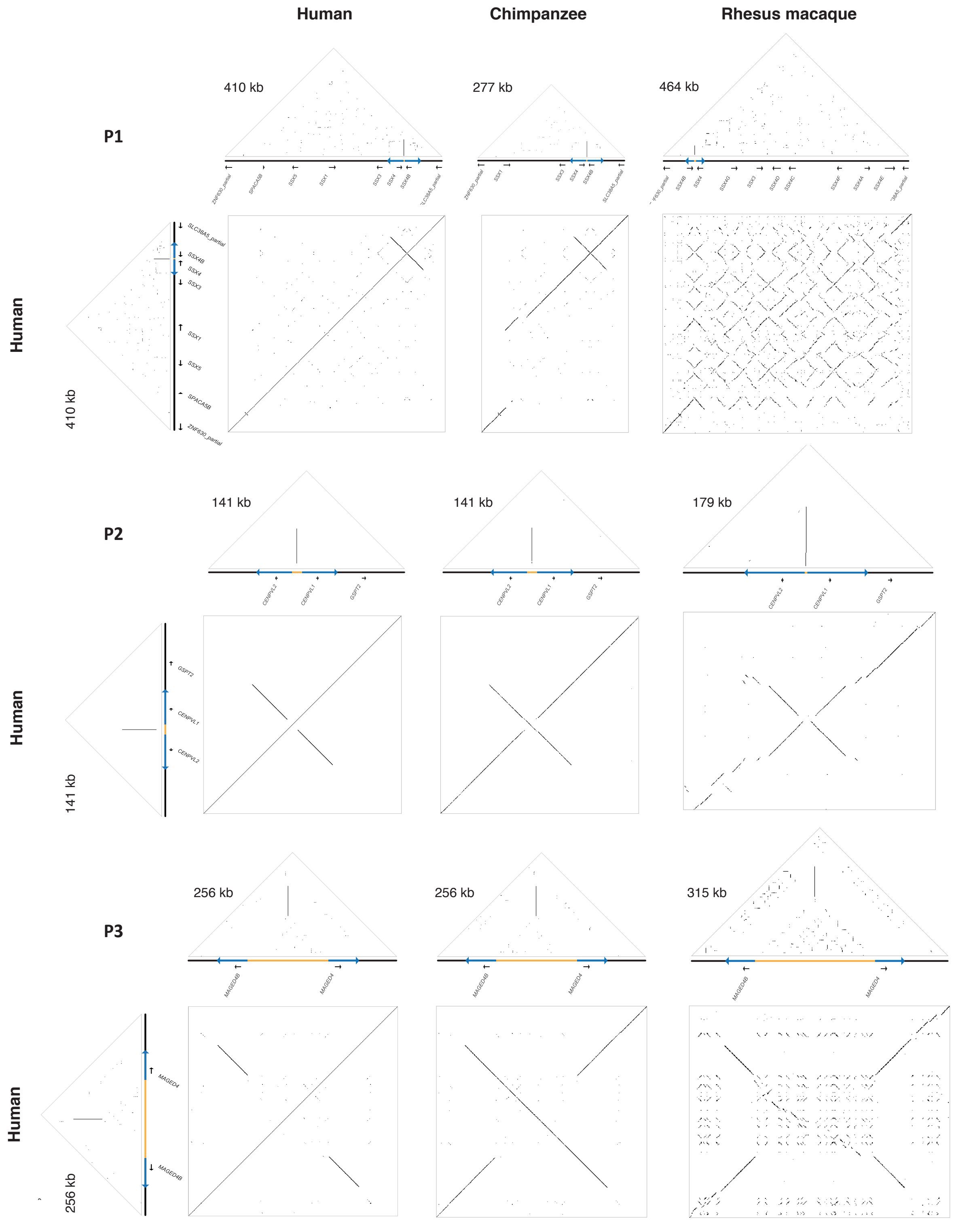


**Supplemental Figure S5.** Annotated square and triangular dot plots of primate X palindromes. w=100 for all triangle plots. w=100 for human vs. human square plots, human vs. chimpanzee square plots; w=40 for human vs. rhesus macaque square plots.

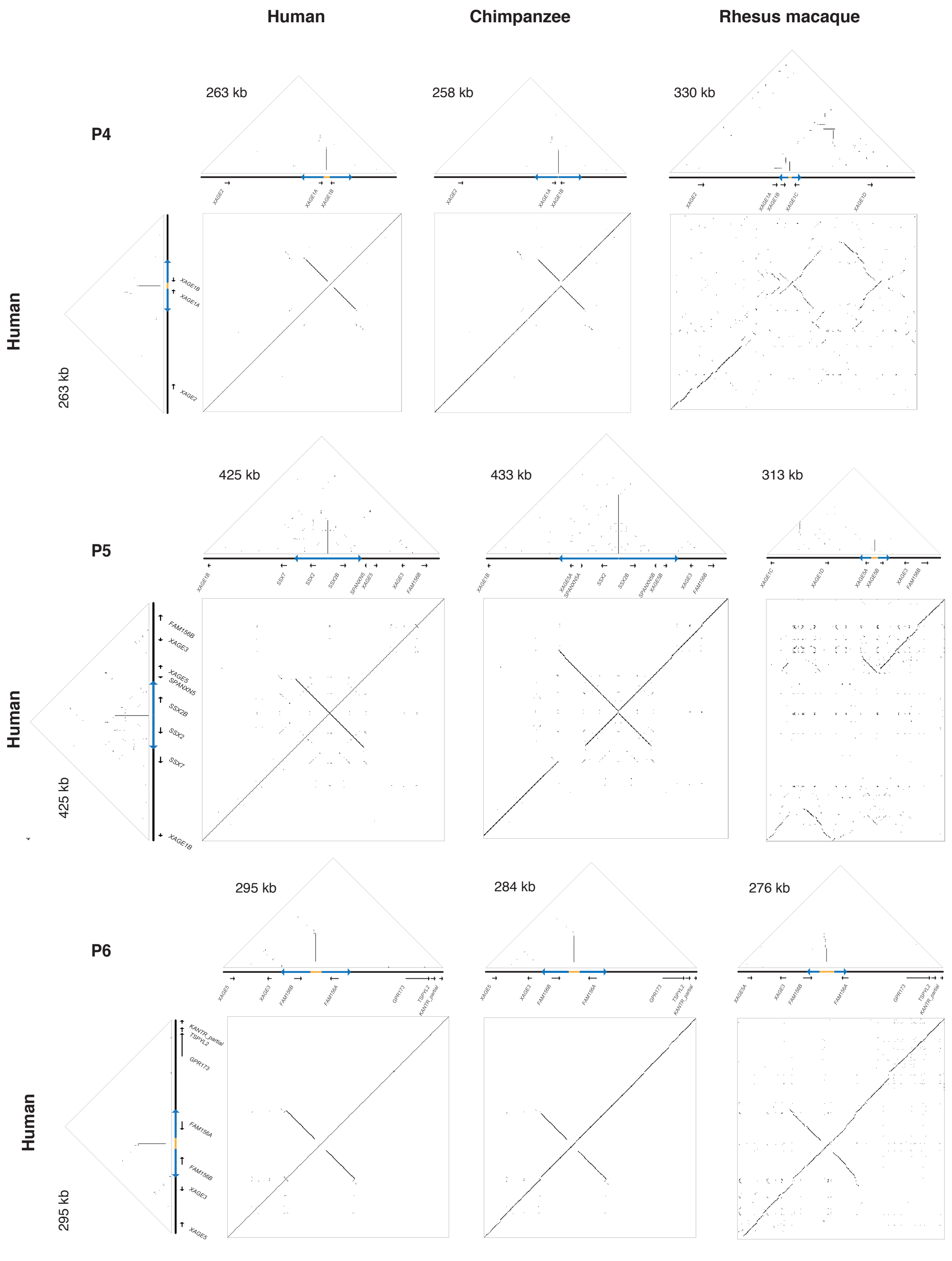


**Supplemental Figure S5.** Annotated square and triangular dot plots of primate X palindromes. w=100 for all triangle plots. w=100 for human vs. human square plots, human vs. chimpanzee square plots; w=40 for human vs. rhesus macaque square plots.

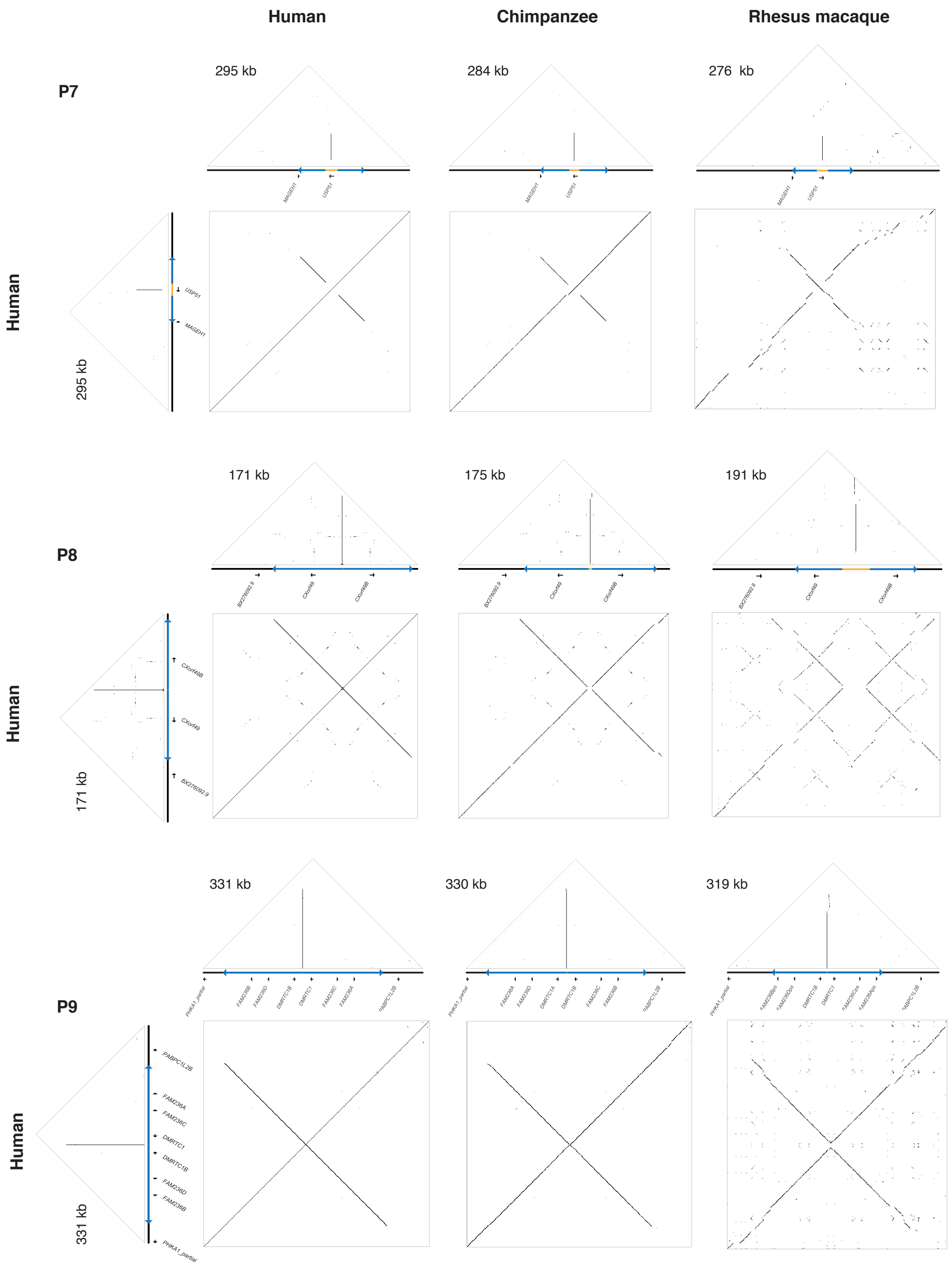


**Supplemental Figure S5.** Annotated square and triangular dot plots of primate X palindromes. w=100 for all triangle plots. w=100 for human vs. human square plots, human vs. chimpanzee square plots; w=40 for human vs. rhesus macaque square plots.

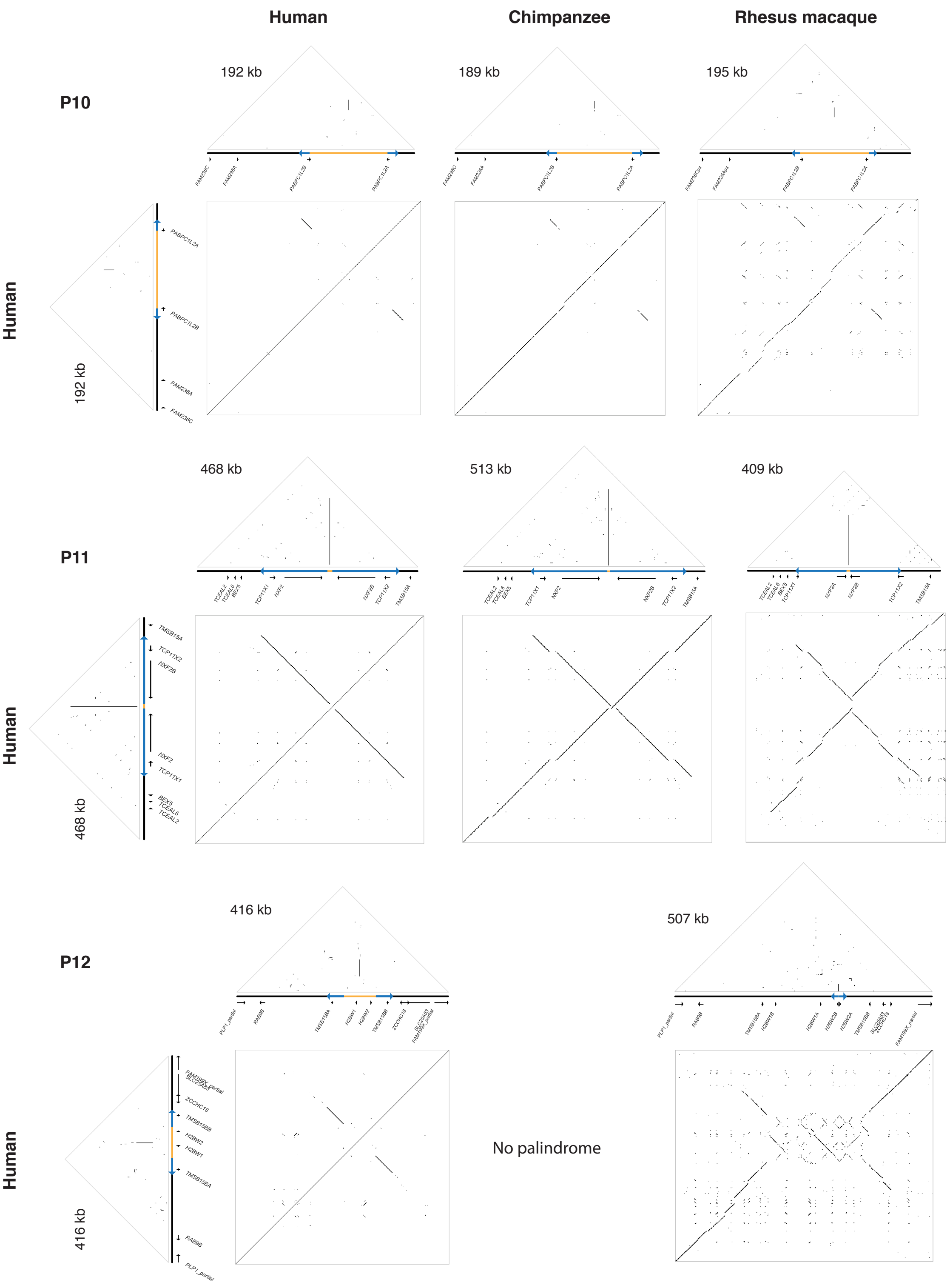


**Supplemental Figure S5.** Annotated square and triangular dot plots of primate X palindromes. w=100 for all triangle plots. w=100 for human vs. human square plots, human vs. chimpanzee square plots; w=40 for human vs. rhesus macaque square plots.

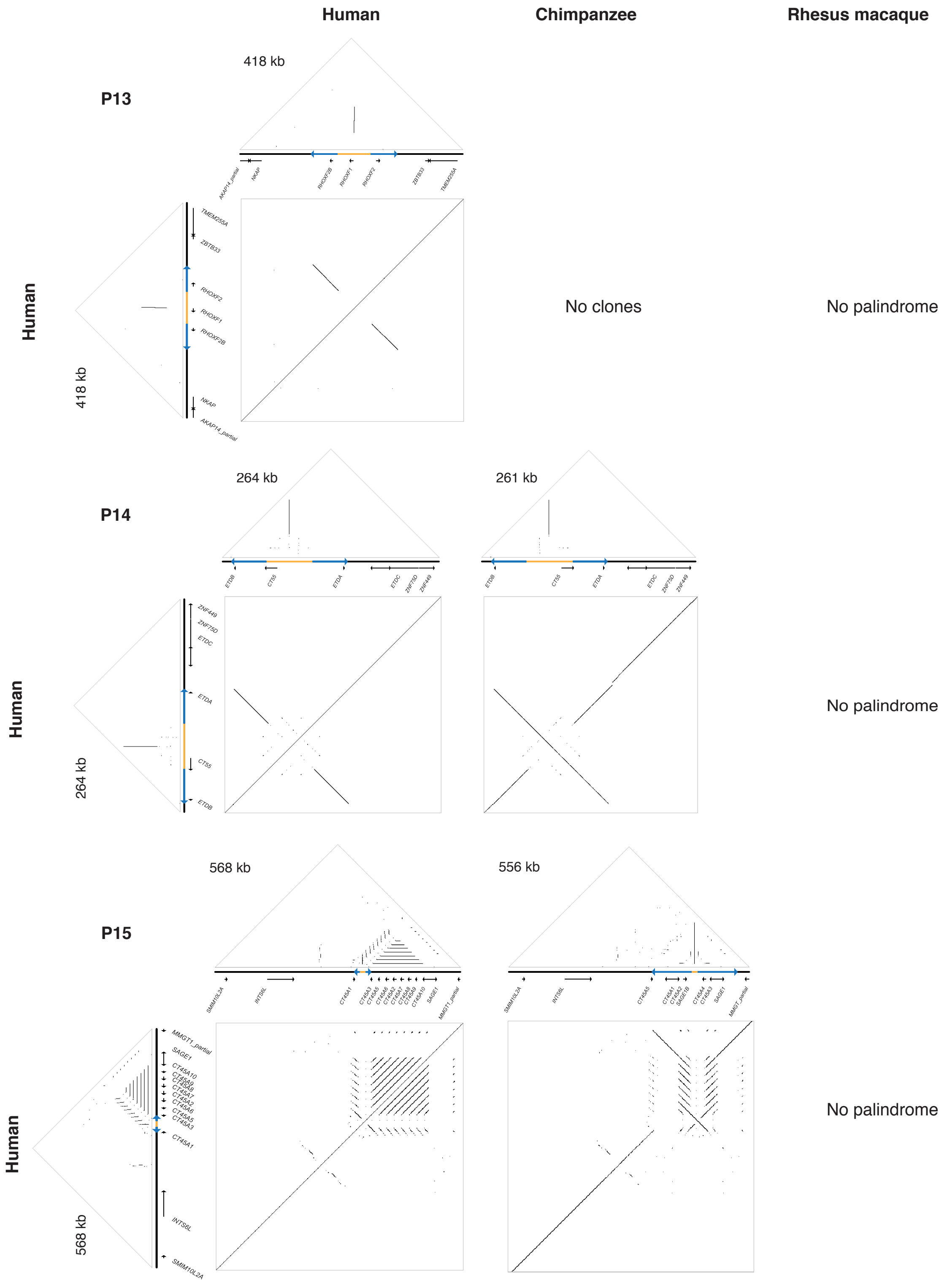


**Supplemental Figure S5.** Annotated square and triangular dot plots of primate X palindromes. w=100 for all triangle plots. w=100 for human vs. human square plots, human vs. chimpanzee square plots; w=40 for human vs. rhesus macaque square plots.

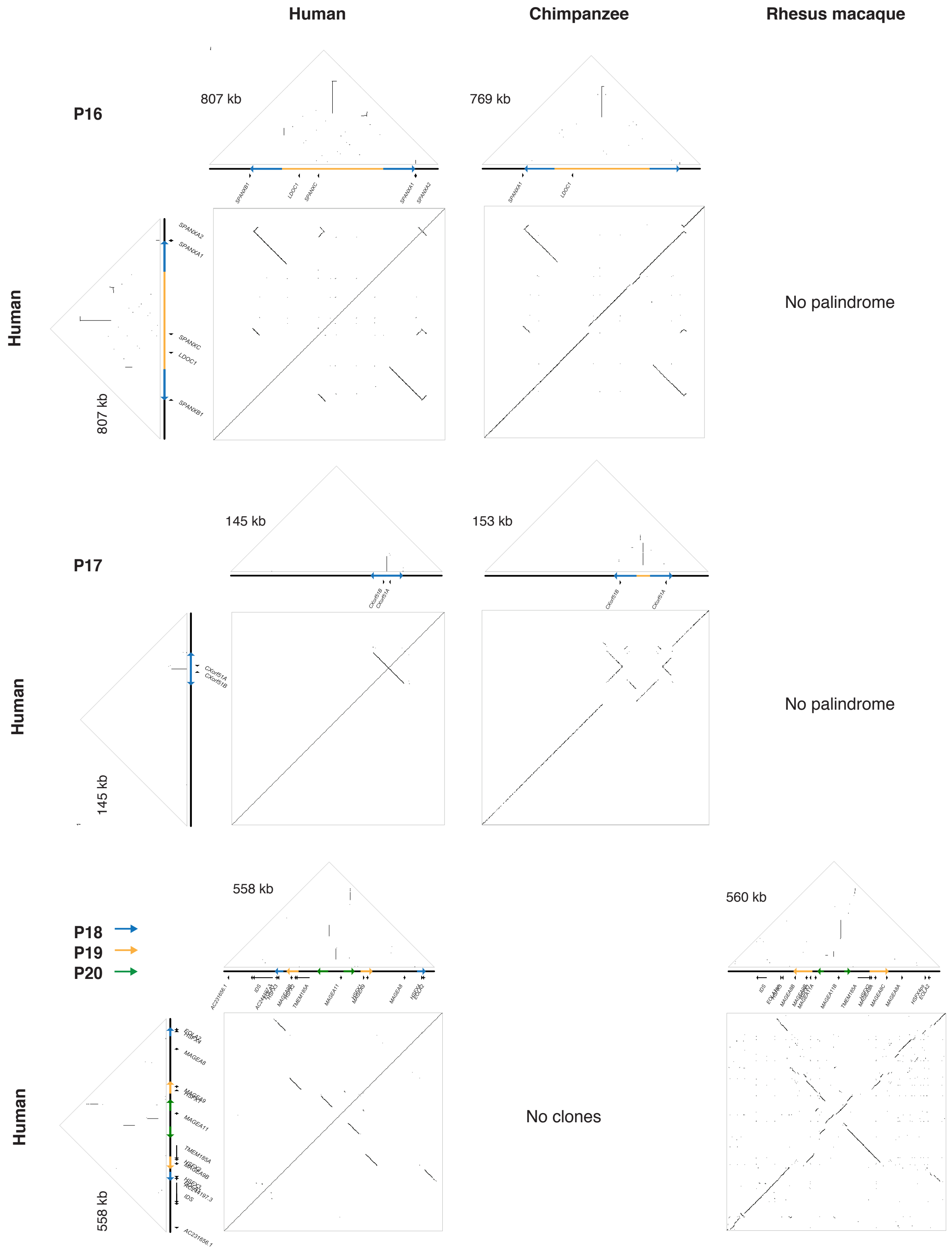


**Supplemental Figure S5.** Annotated square and triangular dot plots of primate X palindromes. w=100 for all triangle plots. w=100 for human vs. human square plots, human vs. chimpanzee square plots; w=40 for human vs. rhesus macaque square plots.

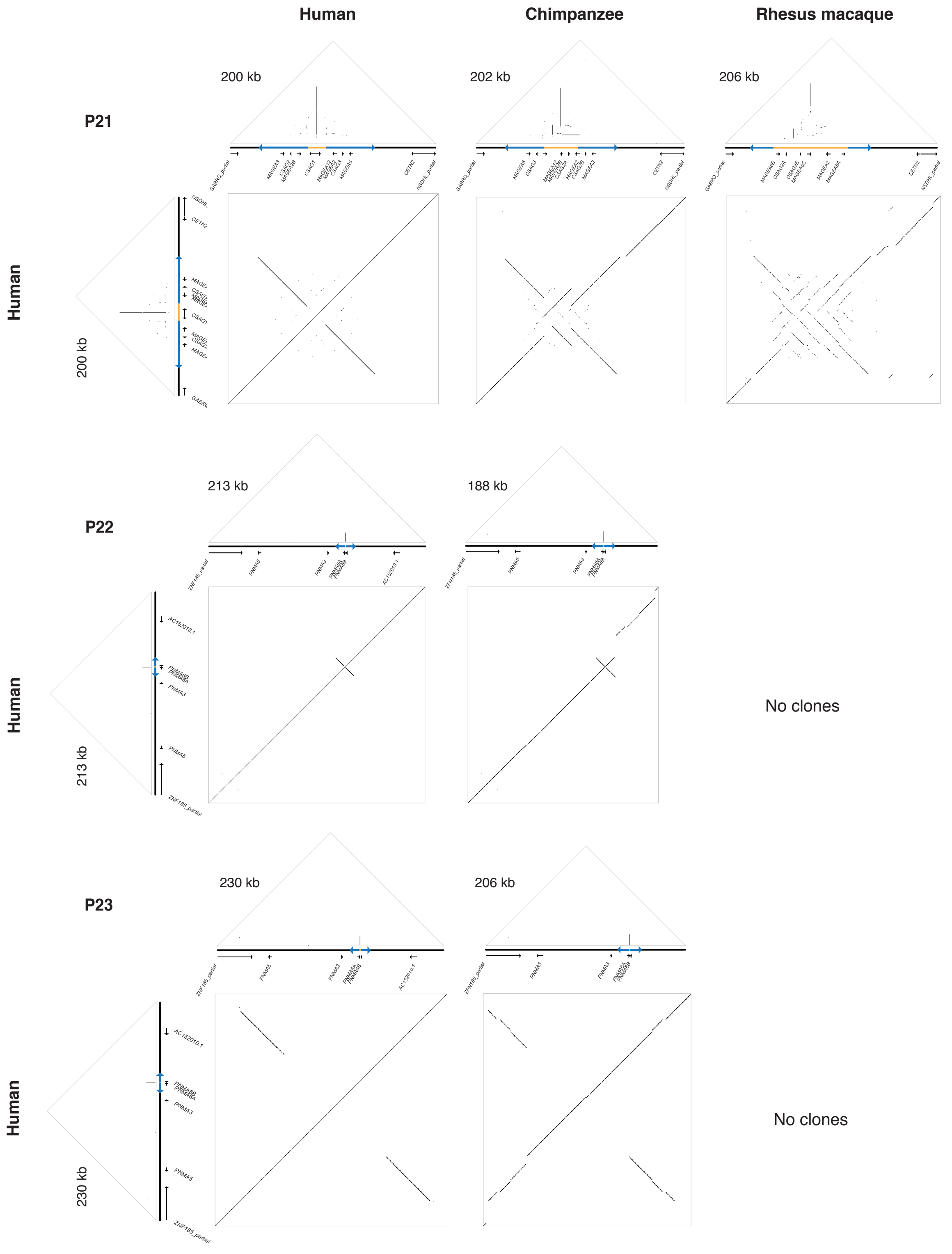


**Supplemental Figure S5.** Annotated square and triangular dot plots of primate X palindromes. w=100 for all triangle plots. w=100 for human vs. human square plots, human vs. chimpanzee square plots; w=40 for human vs. rhesus macaque square plots.

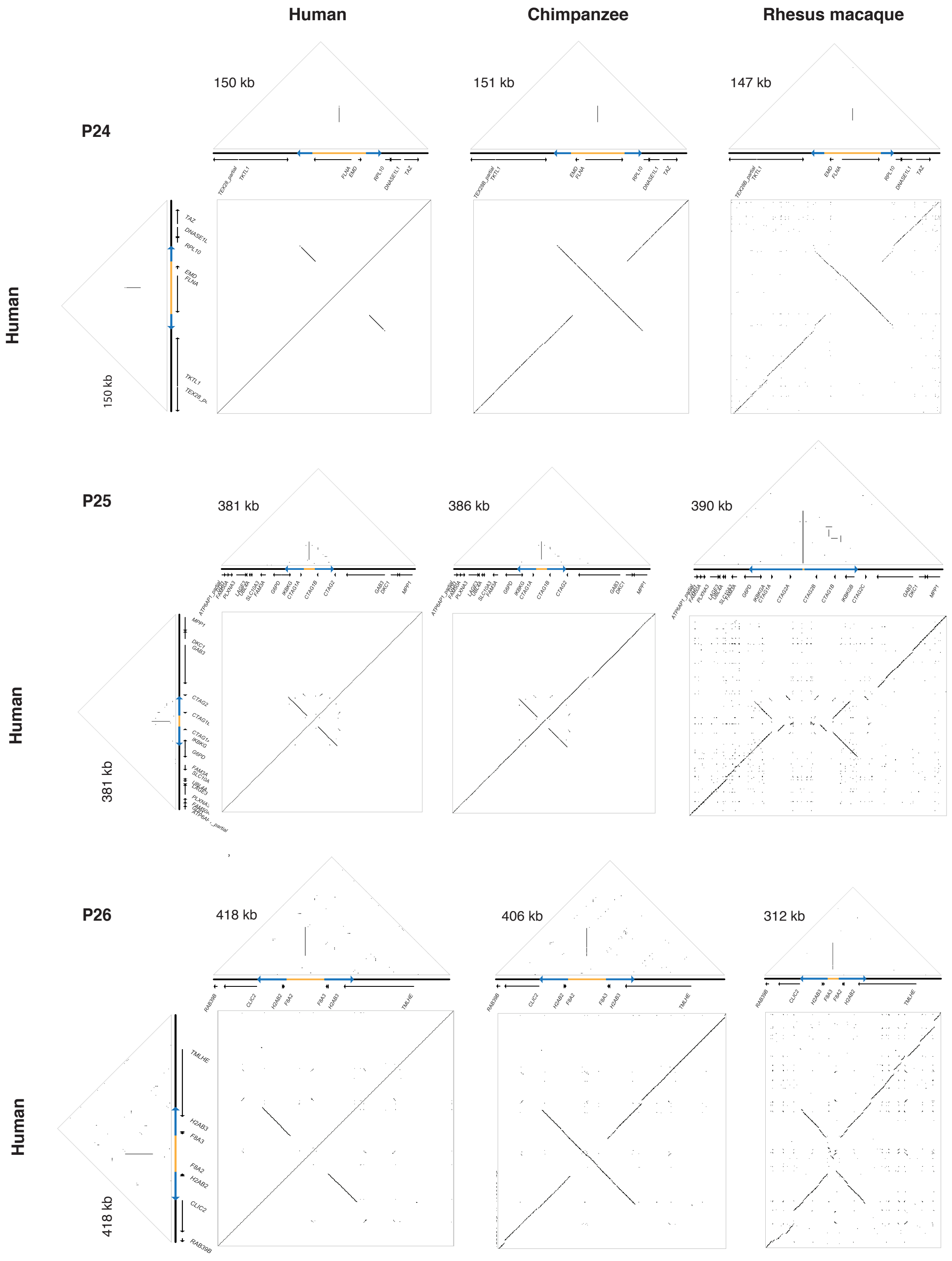


**Supplemental Figure S5.** Annotated square and triangular dot plots of primate X palindromes. w=100 for all triangle plots. w=100 for human vs. human square plots, human vs. chimpanzee square plots; w=40 for human vs. rhesus macaque square plots.

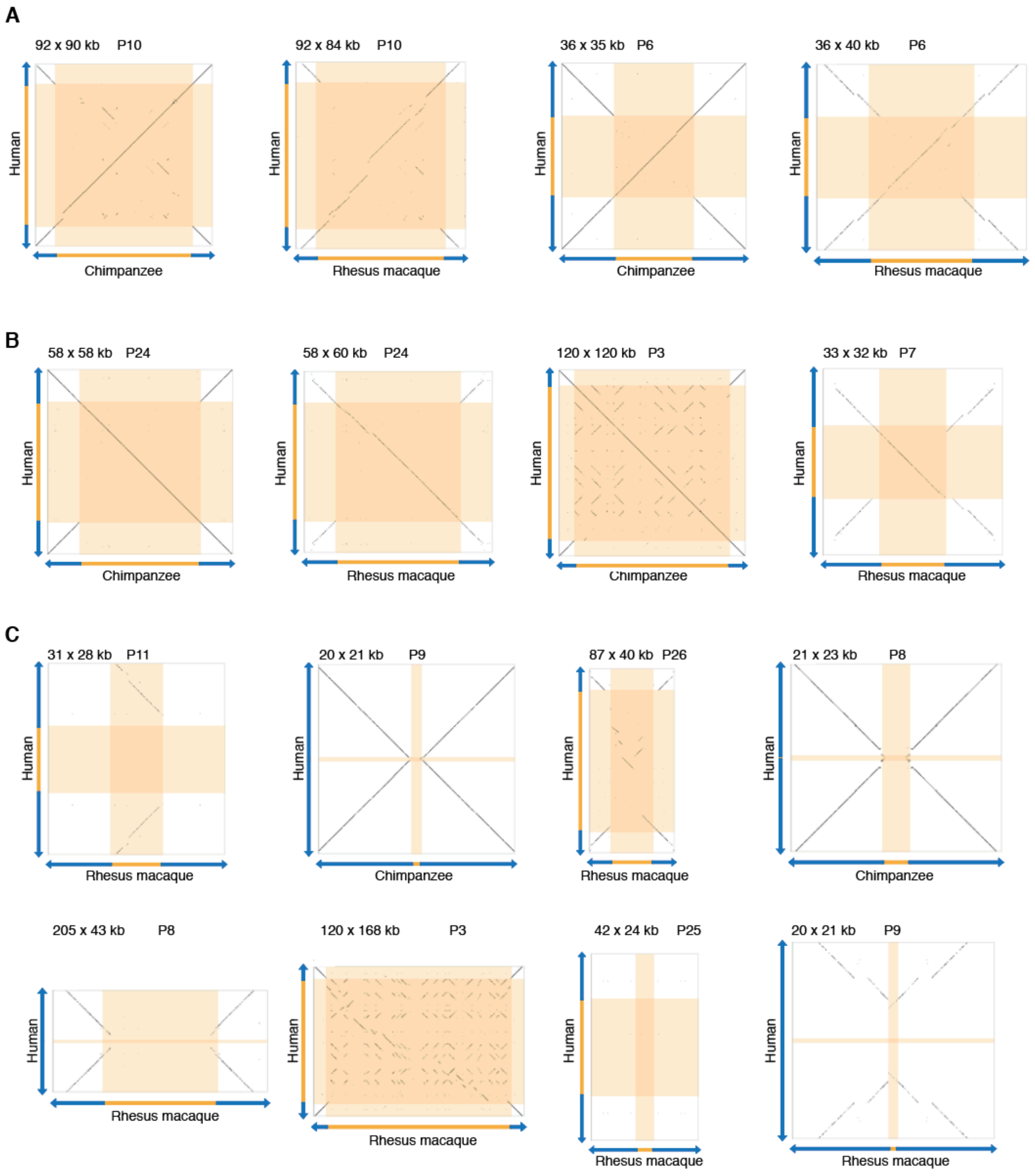


**Supplemental Figure S6**. Additional examples of spacer configurations in orthologous palindromes. All plots show the inner 10 kb of palindrome arms plus the spacer. w=40 for human vs. rhesus macaque comparisons; w=100 for human vs. chimpanzee comparisons. a) Human configuration, b) Inversions, c) Non-orthologous spacers.

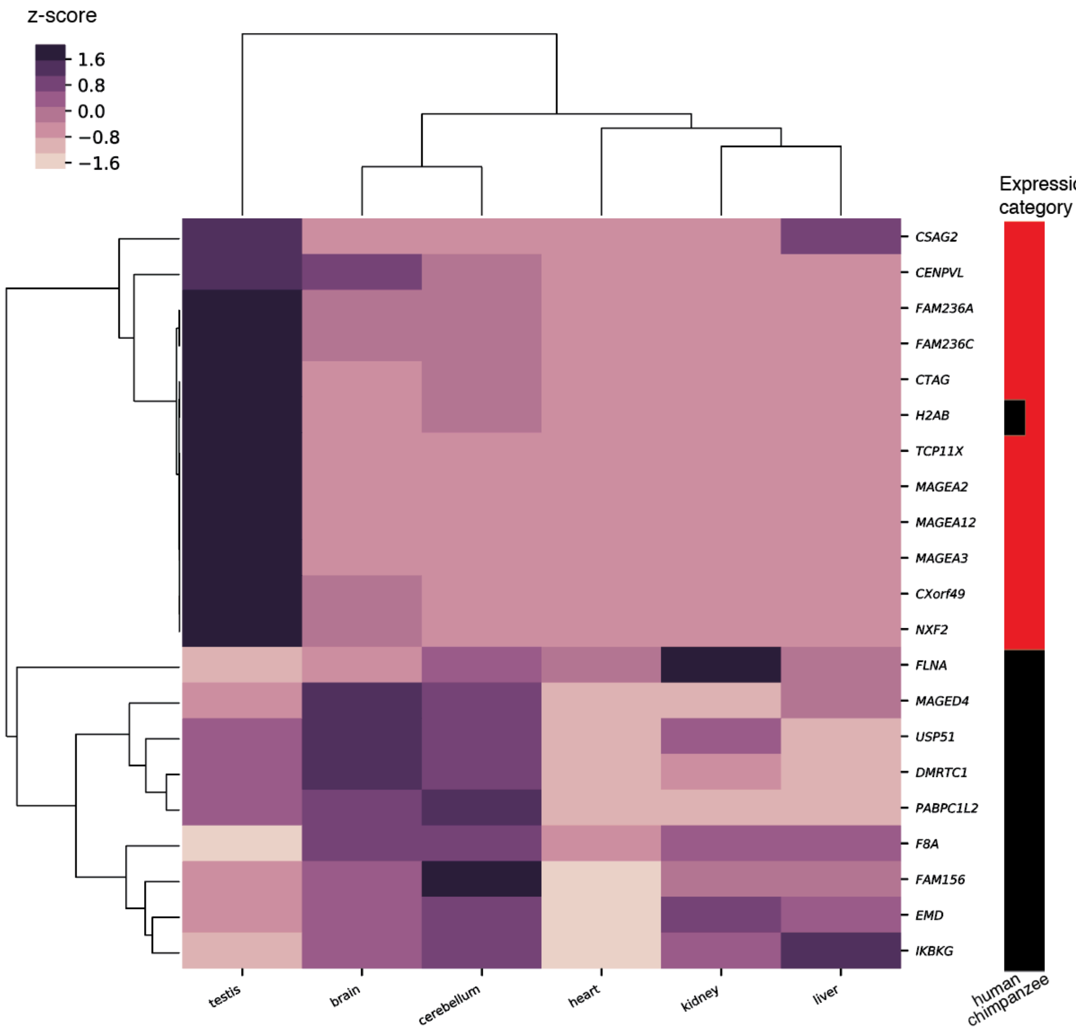


**Supplemental Figure S7.** Expression of gene families from palindromes shared by human, chimpanzee, and macaque in chimpanzee. Data from Brawand et al. 2011 was re-analyzed with kallisto. Each row shows averaged expression from one gene family. Row and column orders were determined by hierarchical clustering. Expression category: Shows whether expression is testis-biased (red) or broad (black) in the indicated species. Testis-biased: Minimum 2 TPM in testis, and testis accounts for >25% of log2 normalized expression summed across all tissues. Broad: All other expressed genes.

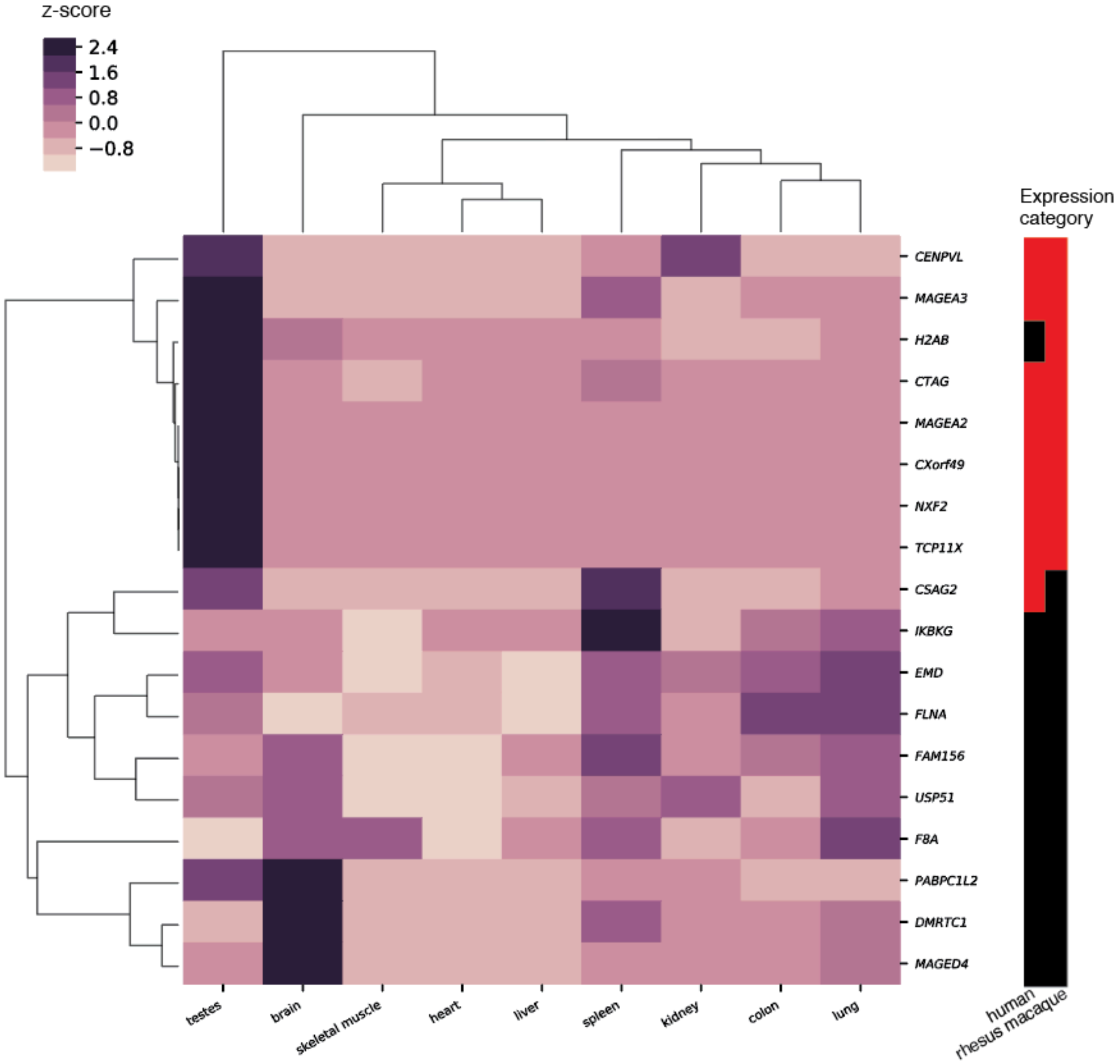


**Supplemental Figure S8.** Expression of gene families from palindromes shared by human, chimpanzee, and macaque in macaque. Data from Merkin et al. 2012 was re-analyzed with kallisto. Each row shows averaged expression from one gene family. Row and column orders were determined by hierarchical clustering. Expression category: Shows whether expression is testis-biased (red) or broad (black) in the indicated species. Testis-biased: Minimum 2 TPM in testis, and testis accounts for >25% of log2 normalized expression summed across all tissues. Broad: All other expressed genes.

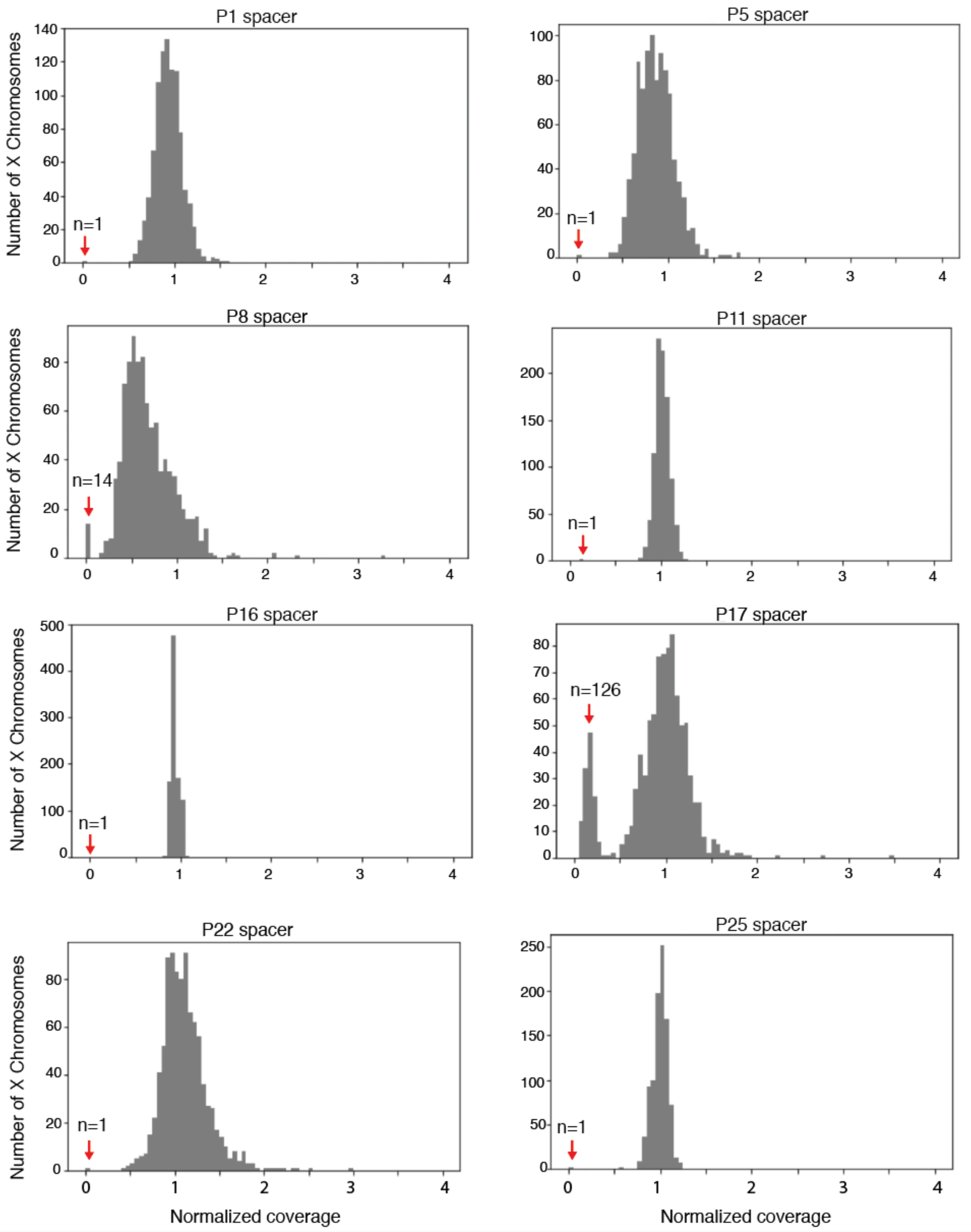


**Supplemental Figure S9**. Normalized coverage depths for eight palindrome spacers with at least one deletion in the 1000 Genomes dataset. Coverage depth for the ninth palindrome with spacer deletions, P2, is shown in Figure 6A.

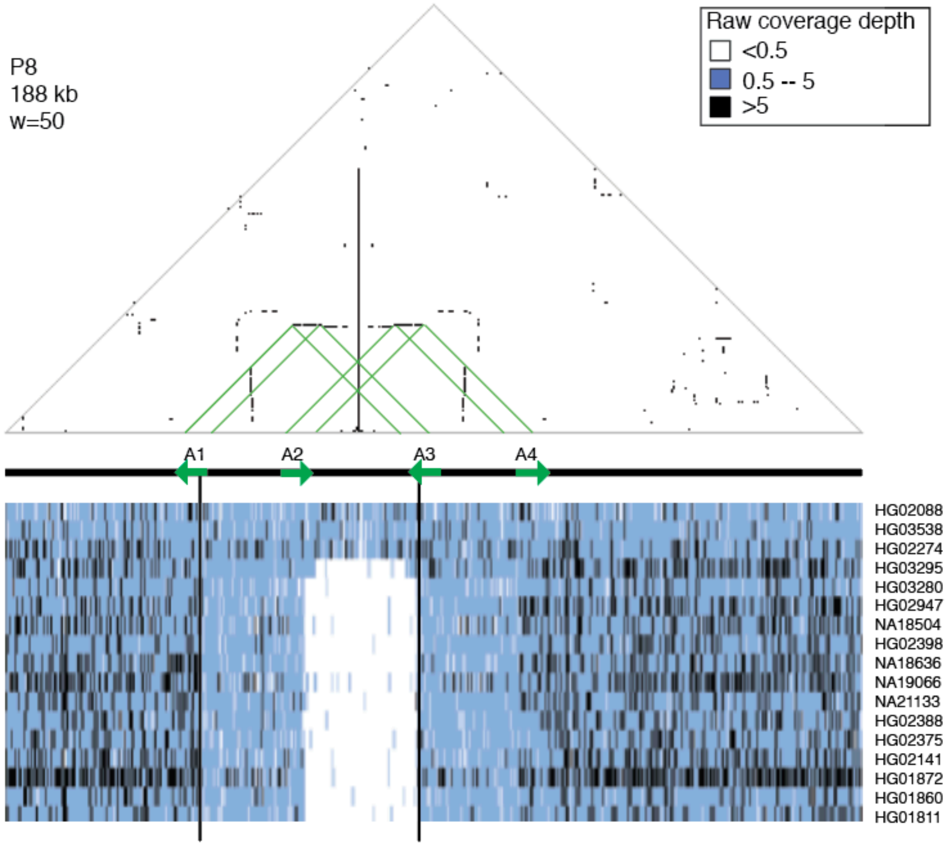


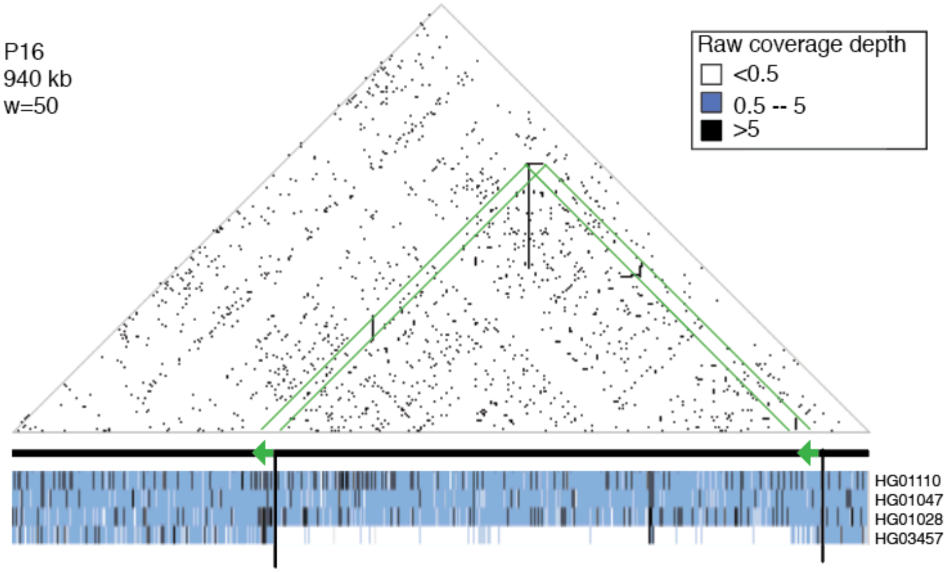


**Supplemental Figure S10.** Human spacer deletions with breakpoints within tandem repeats. Green arrows: Tandem repeats suspected to cause deletions through NAHR. In the case of P8, the deletion could have occurred with equal probability between arrows A1 & A3, or A2 & A4. Note that the suspected P8 deletion spans areas of no coverage (white) and reduced coverage (lighter blue, from approximately A1 to A2); the copy number of the lighter blue region is reduced from 2 to 1.

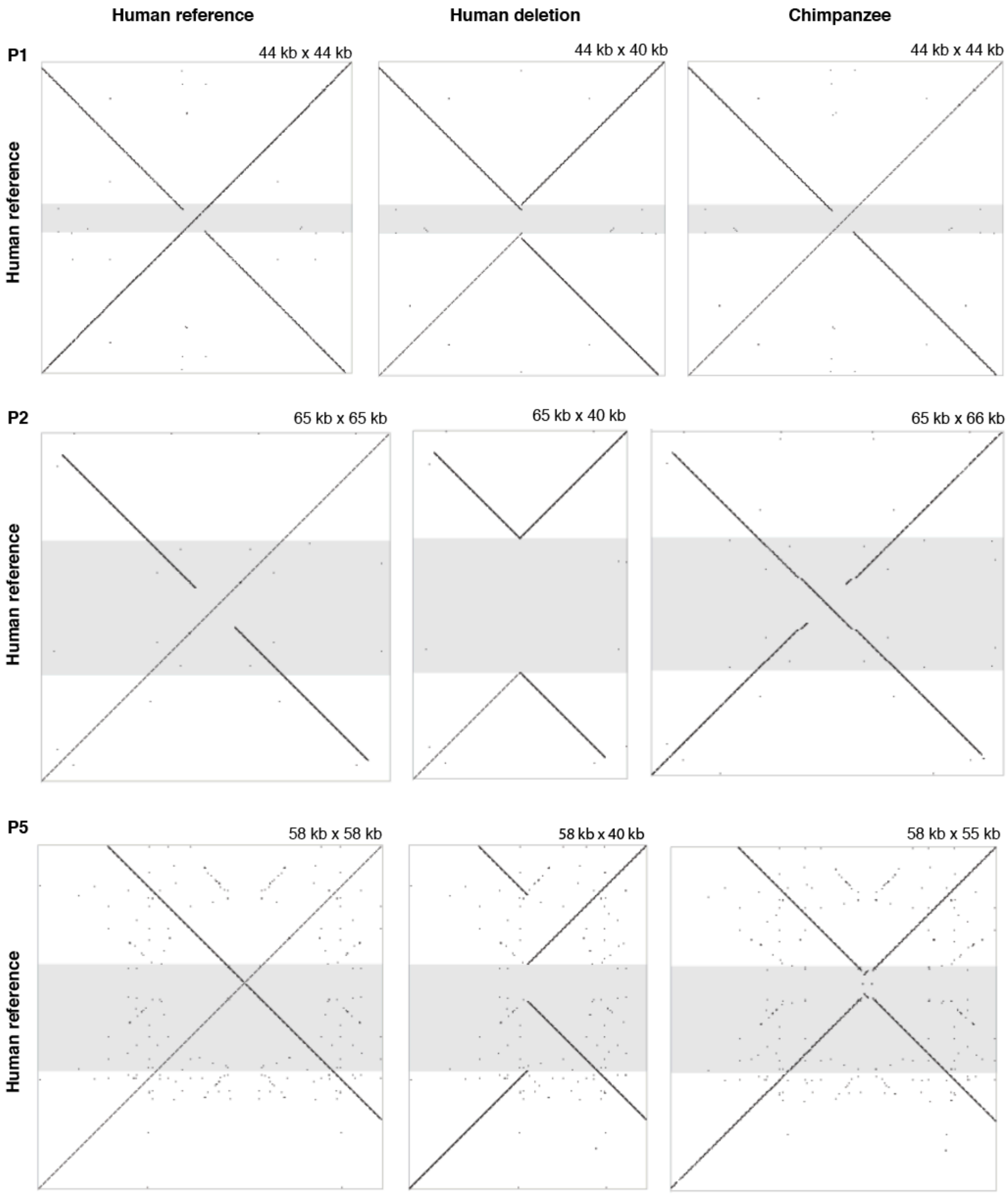


**Supplemental Figure S11.** Structural comparisons between human reference, human deletion, and chimpanzee for nine X palindromes with at least one spacer deletion. w=30 for all square dot plots. Position of the human X spacer deletion is highlighted in gray. For all nine palindromes, most or all of the of sequence absent in the human deletion is present in chimpanzee, confirming that the human structural polymorphism results from deletion rather than insertion.

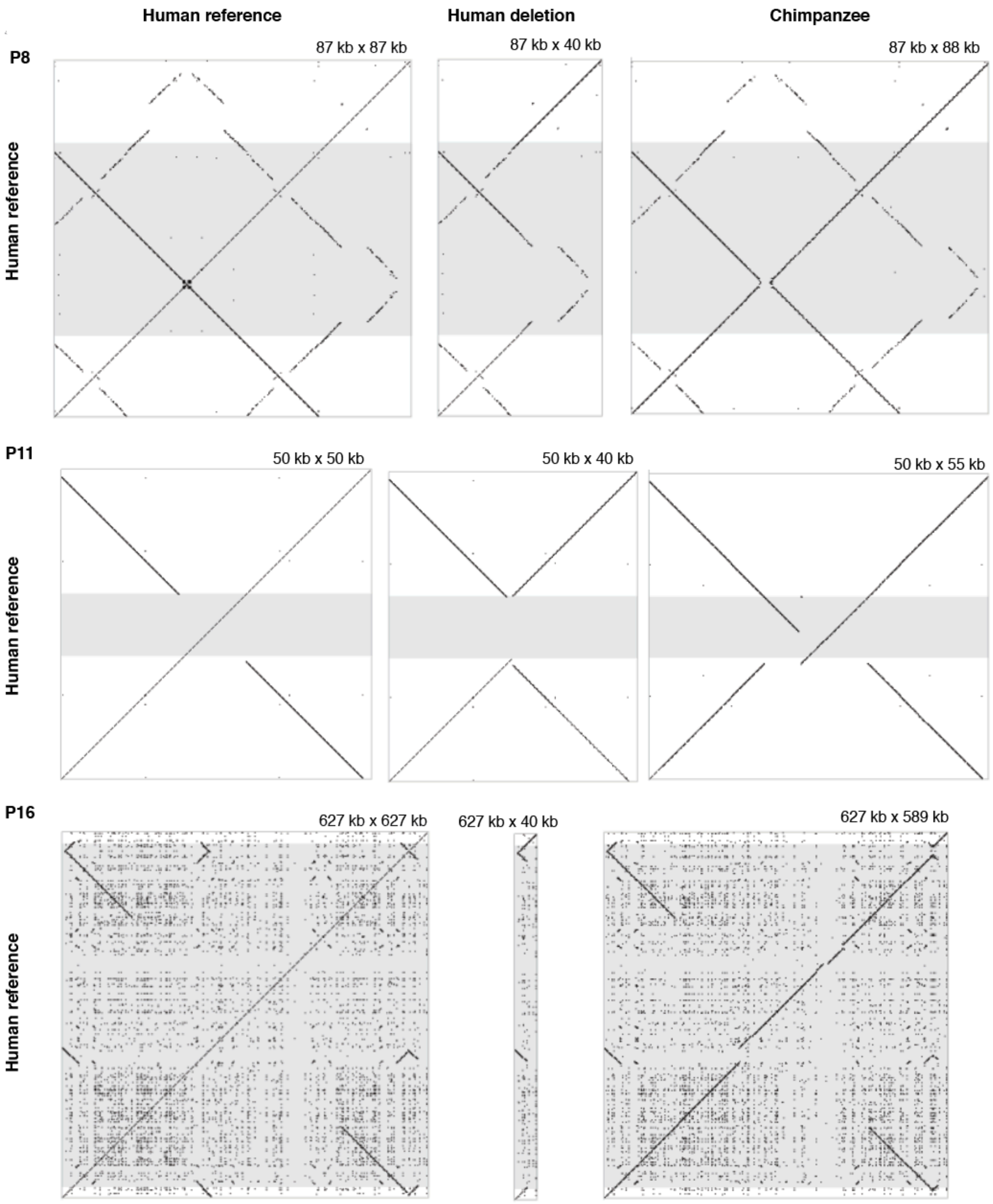


**Supplemental Figure S11.** Structural comparisons between human reference, human deletion, and chimpanzee for nine X palindromes with at least one spacer deletion. w=30 for all square dot plots. Position of the human X spacer deletion is highlighted in gray. For all nine palindromes, most or all of the of sequence absent in the human deletion is present in chimpanzee, confirming that the human structural polymorphism results from deletion rather than insertion.

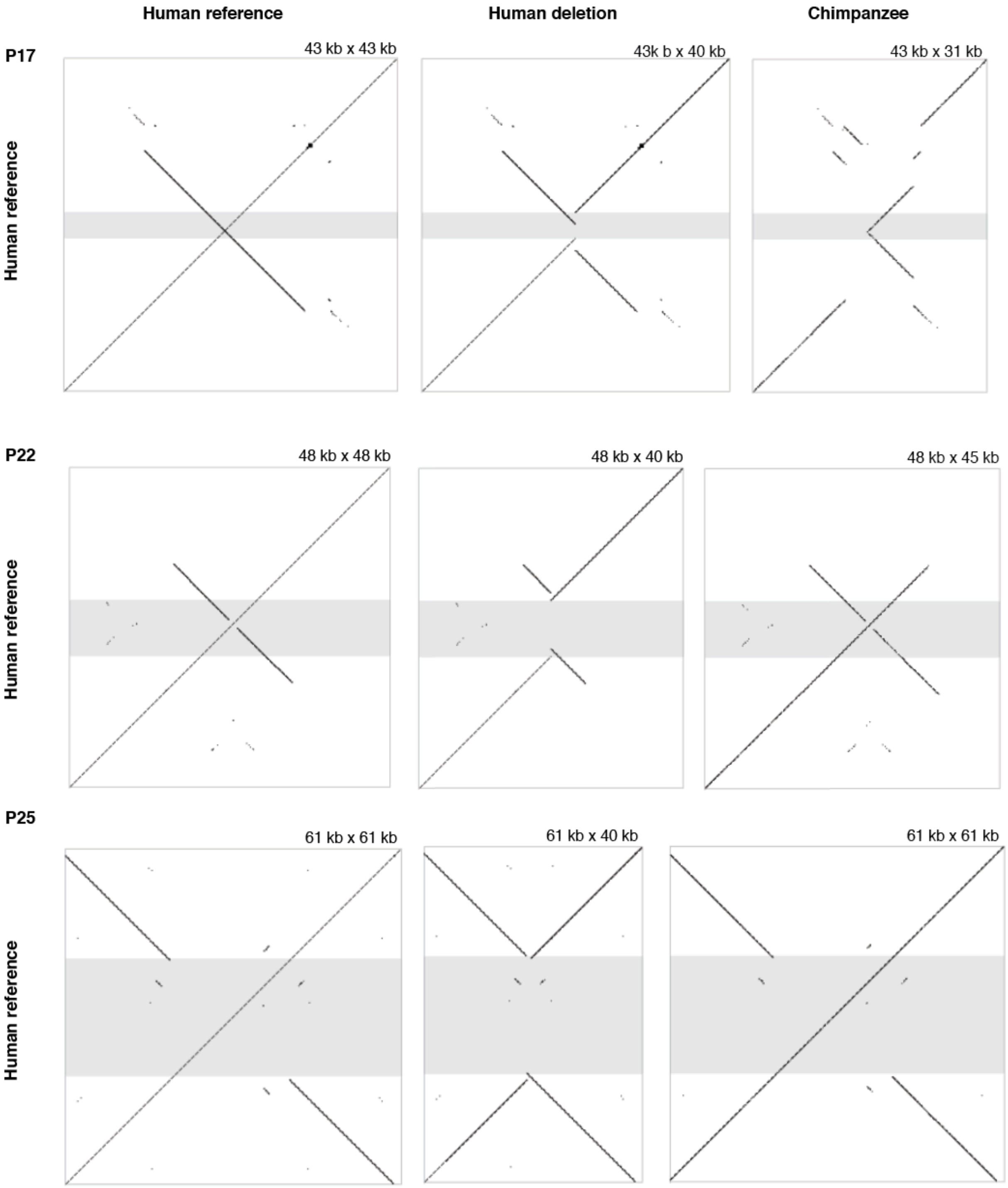


**Supplemental Figure S11.** Structural comparisons between human reference, human deletion, and chimpanzee for nine X palindromes with at least one spacer deletion. w=30 for all square dot plots. Position of the human X spacer deletion is highlighted in gray. For all nine palindromes, most or all of the of sequence absent in the human deletion is present in chimpanzee, confirming that the human structural polymorphism results from deletion rather than insertion.

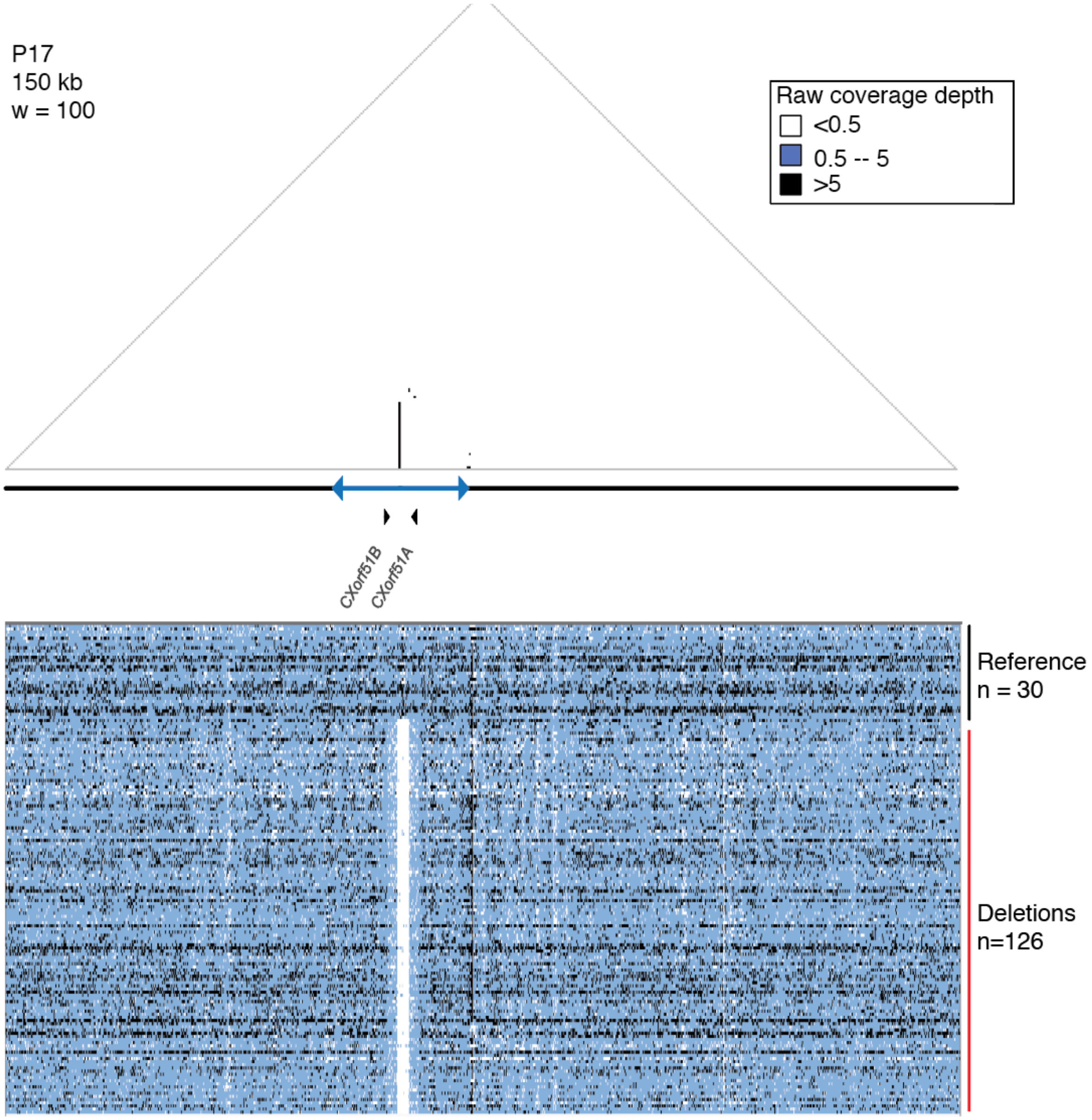


**Supplemental Figure S12**. Coverage depth for X Chromosomes with P17 spacer deletions. Tracks are shown for all 126 X Chromosomes with P17 spacer deletions, plus 30 randomly chosen X Chromosomes with the reference structure.

```
Proximal reference  GCTTTCTGTCACTATCTGTTTCTATTTTCTGGAGGTTTGTAAGAATGGAATCATAAGATATGTACTCTTT
                    ...................................
P17 deletion        GCTTTCTGTCACTATCTGTTTCTATTTTCTGGAGGCTCACTGGTCACCTTTGCCATACTGAATTGTCCC
                                                 ....................................
Distal reference     TCTGTTCGTCCACATCTTCATTAGGCTTCTGTGGCTCACTGGTCACCTTTGCCATACTGAATTGTCCC
```

**Supplemental Figure S13.** Junction for P17 spacer deletion. Proximal reference: chrX: 146811296-146811365. Distal reference: chrX: 146814668-146814736. Two base pairs (GG, purple) overlap between the breakpoints.

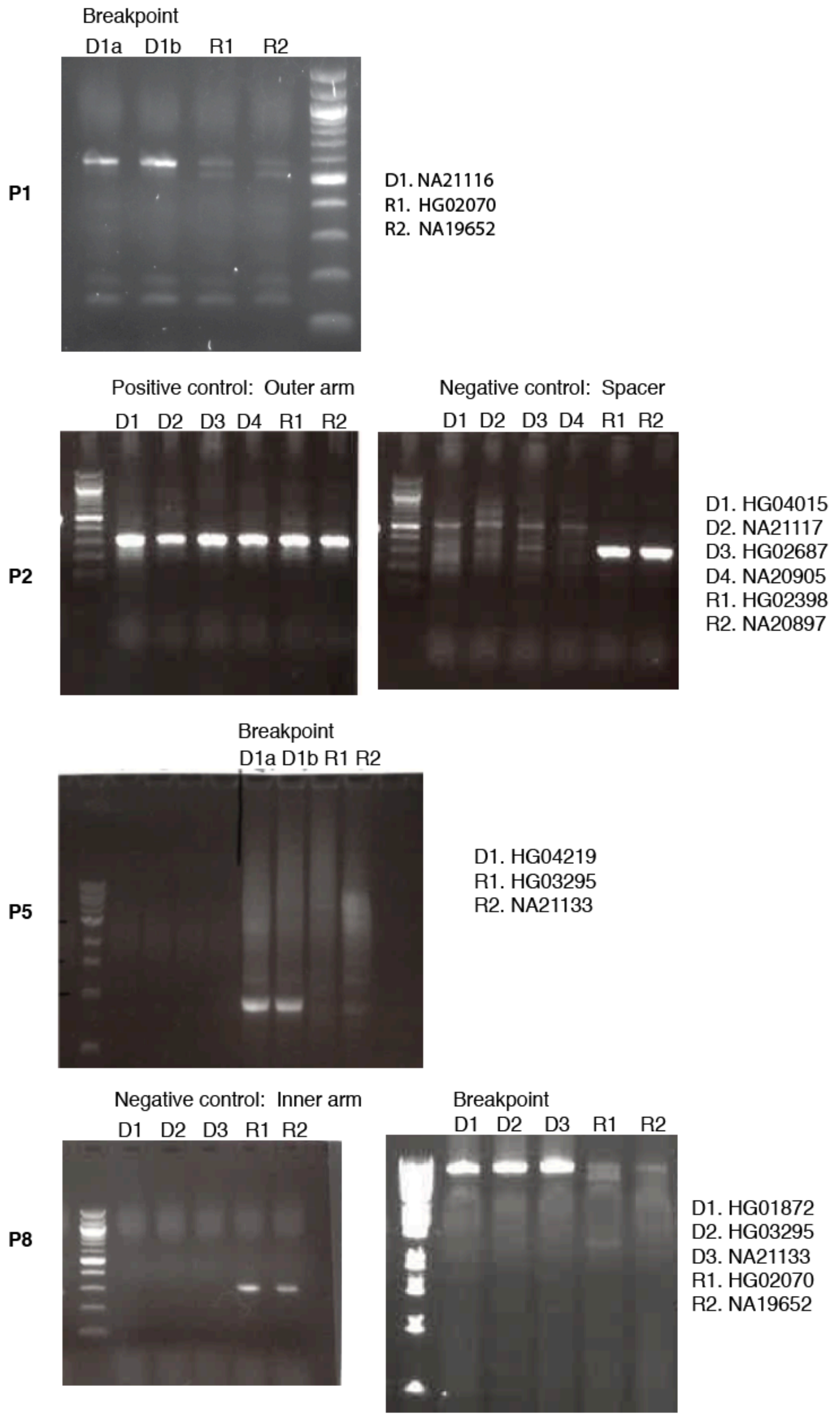


**Supplemental Figure S14.** Verification of human X-palindrome spacer deletions. PCR primers were designed based on deletion breakpoints from split reads or, in cases where reads spanning the breakpoint could not be found, based on the estimated deletion breakpoints from visualization of coverage depth. Positive control: Sequence expected to be present in both reference samples and deletion samples. Negative control: Sequence expected to be present in reference samples, and absent in deletion samples. Breakpoint: Sequence expected to be present in deletion samples, and absent in reference samples. D = deletion sample, R = reference sample.

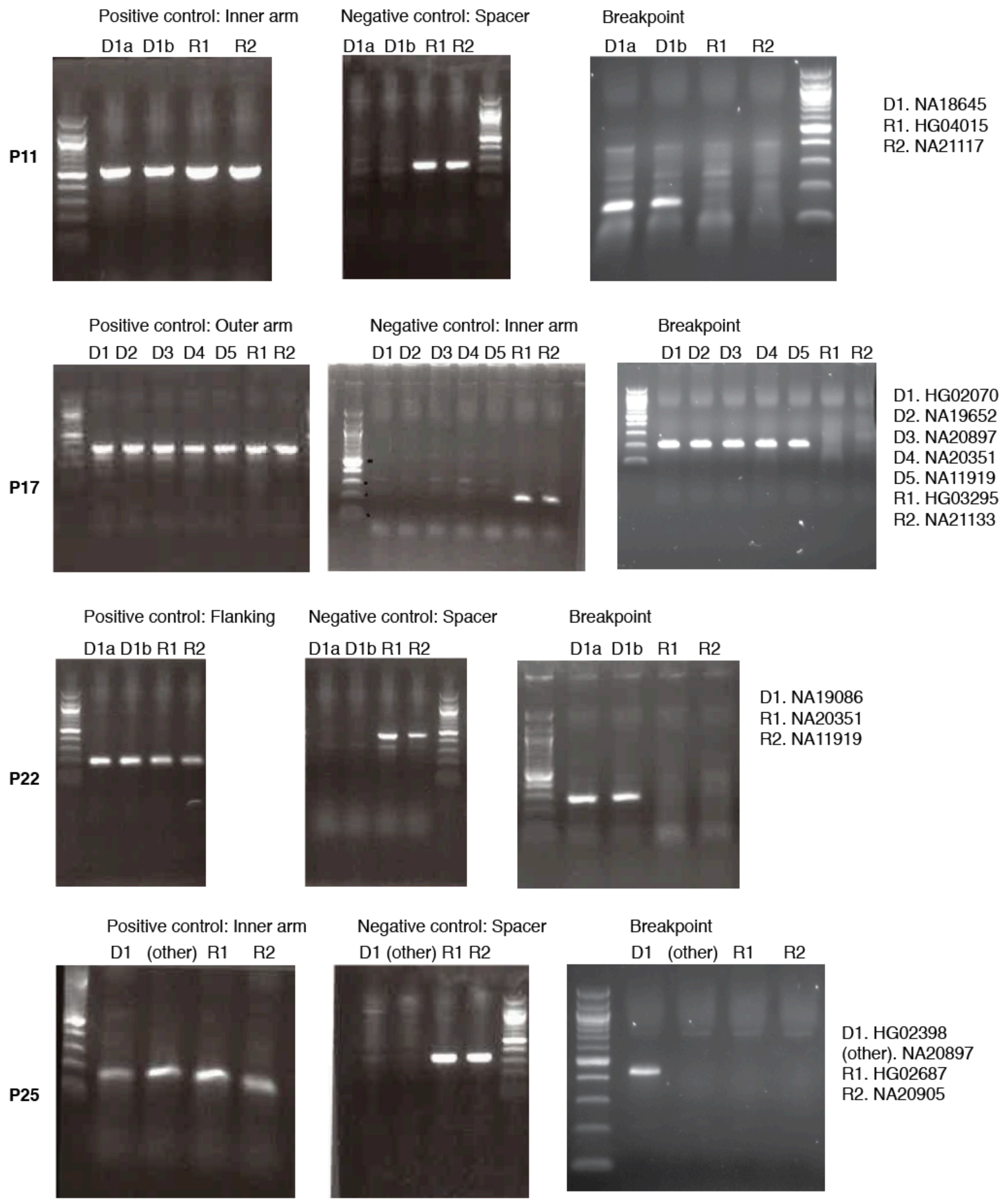


**Supplemental Figure S14.** Verification of human X-palindrome spacer deletions. PCR primers were designed based on deletion breakpoints from split reads or, in cases where reads spanning the breakpoint could not be found, based on the estimated deletion breakpoints from visualization of coverage depth. Positive control: Sequence expected to be present in both reference samples and deletion samples. Negative control: Sequence expected to be present in reference samples, and absent in deletion samples. Breakpoint: Sequence expected to be present in deletion samples, and absent in reference samples. D = deletion sample, R = reference sample.

**Supplemental Table 1:** Coordinates of human X palindromes in hg38. Palindromes were identified using a kmer-based method (see Methods).

| Palindrome | Arm 1 coordinates | Spacer coordinates | Arm 2 coordinates |
|---|---|---|---|
| P1 | chrX: 48367308 - 48396305 | chrX: 48396306 - 48399118 | chrX: 48399119 - 48428123 |
| P2 | chrX: 51668115 - 51692963 | chrX: 51692964 - 51700361 | chrX: 51700362 - 51725226 |
| P3 | chrX: 52040713 - 52077110 | chrX: 52077111 - 52176982 | chrX: 52176983 - 52213380 |
| P4 | chrX: 52473449 - 52502219 | chrX: 52502220 - 52510667 | chrX: 52510668 - 52539437 |
| P5 | chrX: 52670289 - 52728969 | chrX: 52728970 - 52729454 | chrX: 52729455 - 52788214 |
| P6 | chrX: 52882200 - 52920138 | chrX: 52920139 - 52935673 | chrX: 52935674 - 52973611 |
| P7 | chrX: 55453561 - 55480141 | chrX: 55480142 - 55493061 | chrX: 55493062 - 55519639 |
| P8 | chrX: 71683199 - 71740534 | chrX: 71740535 - 71741054 | chrX: 71741055 - 71798363 |
| P9 | chrX: 72741066 - 72860190 | chrX: 72860191 - 72860605 | chrX: 72860606 - 72979767 |
| P10 | chrX: 72996086 - 73005278 | chrX: 73005279 - 73077748 | chrX: 73077749 - 73086935 |
| P11 | chrX: 102197589 - 102338170 | chrX: 102338171 - 102348932 | chrX: 102348933 - 102489526 |
| P12 | chrX: 103955950 - 103987907 | chrX: 103987908 - 104050879 | chrX: 104050880 - 104082911 |
| P13 | chrX: 120038201 - 120086776 | chrX: 120086777 - 120149524 | chrX: 120149525 - 120198163 |
| P14 | chrX: 135116122 - 135158129 | chrX: 135158130 - 135214412 | chrX: 135214413 - 135256414 |
| P15 | chrX: 135723708 - 135734732 | chrX: 135734733 - 135748584 | chrX: 135748585 - 135759685 |
| P16 | chrX: 141005250 - 141114518 | chrX: 141114519 - 141472891 | chrX: 141472892 - 141582141 |
| P17 | chrX: 146801856 - 146812219 | chrX: 146812220 - 146812295 | chrX: 146812296 - 146822659 |
| P18 | chrX: 149542167 - 149562176 | chrX: 149562177 - 149917319 | chrX: 149917320 - 149937339 |
| P19 | chrX: 149573130 - 149602230 | chrX: 149602231 - 149767142 | chrX: 149767143 - 149796257 |
| P20 | chrX: 149654519 - 149681126 | chrX: 149681127 - 149722142 | chrX: 149722143 - 149748749 |
| P21 | chrX: 152678580 - 152725390 | chrX: 152725391 - 152743079 | chrX: 152743080 - 152789894 |
| P22 | chrX: 153066010 - 153074417 | chrX: 153074418 - 153075611 | chrX: 153075612 - 153084043 |
| P23 | chrX: 153106025 - 153149489 | chrX: 153149490 - 153250484 | chrX: 153250485 - 153293919 |
| P24 | chrX: 154337197 - 154347246 | chrX: 154347247 - 154384865 | chrX: 154384866 - 154394951 |
| P25 | chrX: 154555880 - 154591327 | chrX: 154591328 - 154613094 | chrX: 154613095 - 154648556 |
| P26 | chrX: 155336691 - 155386727 | chrX: 155386728 - 155453980 | chrX: 155453981 - 155504550 |

**Supplemental Table 2:** Clones sequenced for this project. Some clones contained portions of multiple palindromes. "Finished": Clone sequence was supported by complete Illumina coverage; "Prefinished": Clone had at least one stretch of sequence supported by one or more nanopore reads, but no Illumina reads.

| Clone | Palindrome | Status | # full length reads |
|---|---|---|---|
| CH250-106M20 | P7 | Prefinished | 5 |
| CH250-114J18 | P9/P10 | Prefinished | 0 |
| CH250-119L11 | P16 | Finished | 0 |
| CH250-120L20 | P18/P19/P20 | Prefinished | 29 |
| CH250-136N6 | P3 | Finished | 12 |
| CH250-137I15 | P26 | Prefinished | 8 |
| CH250-138B21 | P12 | Prefinished | 2 |
| CH250-149O24 | P1 | Prefinished | 8 |
| CH250-150I6 | P15 | Prefinished | 0 |
| CH250-163K20 | P18/P19/P20 | Prefinished | 0 |
| CH250-168E3 | P21 | Prefinished | 3 |
| CH250-174F12 | P21 | Prefinished | 19 |
| CH250-184A21 | P18/P19/P20 | Prefinished | 3 |
| CH250-191K20 | P2 | Finished | 13 |
| CH250-197O3 | P6 | Prefinished | 6 |
| CH250-214O8 | P17 | Finished | 79 |
| CH250-228D11 | P11 | Prefinished | 3 |
| CH250-234D7 | P16 | Finished | 0 |
| CH250-236O7 | P8 | Prefinished | 1 |
| CH250-240H14 | P14 | Finished | 6 |
| CH250-257F3 | P1 | Finished | 4 |
| CH250-257M3 | P1 | Prefinished | 1 |
| CH250-25I12 | P13 | Prefinished | 3 |
| CH250-273C12 | P15 | Prefinished | 2 |
| CH250-280C5 | P16 | Prefinished | 6 |
| CH250-300J22 | P4/P5 | Prefinished | 20 |
| CH250-312L23 | P16 | Finished | 0 |
| CH250-313D10 | P13 | Prefinished | 11 |
| CH250-318K15 | P16 | Finished | 0 |
| CH250-371L16 | P9 | Prefinished | 4 |
| CH250-396M7 | P1 | Prefinished | 3 |
| CH250-397P11 | P15 | Prefinished | 23 |
| CH250-398K19 | P21 | Prefinished | 5 |
| CH250-412K19 | P5/P6 | Prefinished | 16 |

| | | | |
|---|---|---|---|
| CH250-417G7 | P26 | Prefinished | 2 |
| CH250-420A18 | P14 | Finished | 15 |
| CH250-424H13 | P25 | Prefinished | 23 |
| CH250-436M9 | P4/P5 | Prefinished | 0 |
| CH250-440K2 | P13 | Prefinished | 0 |
| CH250-462M8 | P21 | Prefinished | 4 |
| CH250-486E21 | P1 | Prefinished | 0 |
| CH250-487N16 | P26 | Finished | 2 |
| CH250-491H11 | P24 | Prefinished | 2 |
| CH250-493M11 | P25 | Prefinished | 2 |
| CH250-498I16 | P11 | Prefinished | 8 |
| CH250-499B10 | P1 | Finished | 8 |
| CH250-503C21 | P25 | Prefinished | 10 |
| CH250-503N19 | P25 | Prefinished | 2 |
| CH250-504P11 | P18/P19/P20 | Prefinished | 5 |
| CH250-516N14 | P12 | Prefinished | 0 |
| CH250-530N5 | P4/P5 | Finished | 5 |
| CH250-540J3 | P11 | Prefinished | 10 |
| CH250-541H5 | P3 | Prefinished | 0 |
| CH250-547J16 | P18/P19/P20 | Prefinished | 4 |
| CH250-563M7 | P3 | Prefinished | 22 |
| CH250-57C9 | P16 | Finished | 0 |
| CH250-80G22 | P9 | Prefinished | 6 |
| CH250-87B7 | P7 | Prefinished | 7 |
| CH250-92B13 | P12 | Prefinished | 1 |
| CH250-94G2 | P3 | Prefinished | 4 |
| CH250-95D17 | P1 | Prefinished | 2 |
| CH251-130O9 | P4/P5 | Finished | 0 |
| CH251-160A4 | P16 | Prefinished | 7 |
| CH251-161L14 | P7 | Prefinished | 2 |
| CH251-172F20 | P16 | Finished | 0 |
| CH251-177B21 | P2 | Prefinished | 21 |
| CH251-183G21 | P8 | Prefinished | 0 |
| CH251-189G13 | P16 | Prefinished | 0 |
| CH251-239P10 | P26 | Prefinished | 26 |
| CH251-240O17 | P5/P6 | Prefinished | 61 |
| CH251-261H21 | P15 | Prefinished | 4 |
| CH251-277H18 | P7 | Prefinished | 33 |
| CH251-285D14 | P26 | Prefinished | 1 |
| CH251-292E19 | P22/P23 | Prefinished | 8 |

| | | | |
|---|---|---|---|
| CH251-316L7 | P17 | Prefinished | 4 |
| CH251-346A10 | P25 | Prefinished | 12 |
| CH251-34N14 | P4/P5 | Prefinished | 3 |
| CH251-385I8 | P1 | Prefinished | 4 |
| CH251-389B7 | P14 | Prefinished | 20 |
| CH251-397P16 | P3 | Prefinished | 2 |
| CH251-4M24 | P11 | Finished | 21 |
| CH251-504H5 | P10 | Finished | 0 |
| CH251-506D4 | P6 | Prefinished | 0 |
| CH251-50L15 | P15 | Prefinished | 5 |
| CH251-514B7 | P8 | Prefinished | 3 |
| CH251-542A6 | P10 | Prefinished | 12 |
| CH251-542D16 | P12 | Finished | 17 |
| CH251-542E16 | P22 | Prefinished | 4 |
| CH251-550E20 | P12 | Finished | 0 |
| CH251-565G15 | P21 | Prefinished | 3 |
| CH251-571K4 | P15 | Finished | 23 |
| CH251-58J24 | P16 | Finished | 0 |
| CH251-635P13 | P11 | Prefinished | 15 |
| CH251-639F23 | P15 | Prefinished | 3 |
| CH251-64D22 | P16 | Prefinished | 4 |
| CH251-651H9 | P11 | Prefinished | 16 |
| CH251-654E24 | P16 | Prefinished | 25 |
| CH251-657L4 | P9/P10 | Prefinished | 17 |
| CH251-658J15 | P8 | Prefinished | 7 |
| CH251-65E21 | P24 | Prefinished | 1 |
| CH251-671I19 | P3 | Prefinished | 5 |
| CH251-673E12 | P16 | Prefinished | 2 |
| CH251-677L24 | P24/P25 | Prefinished | 8 |
| CH251-702N4 | P16 | Finished | 0 |
| CH251-737G9 | P9/P10 | Prefinished | 1 |
| CH251-73C22 | P2 | Prefinished | 0 |
| CH251-83H5 | P4 | Finished | 0 |

**Supplemental Table S3:** Status of palindromes identified in SHIMS 3.0 in other chimpanzee X assemblies. Missing: No palindrome present in non-SHIMS 3.0 assembly; Incomplete: Part of palindrome present in non-SHIMS 3.0 assembly; Accurate: Full palindrome present in non-SHIMS 3.0 assembly.

| **Palindrome** | **Pan_tro_3.0** | **Clint_PTRv2** | **Arm length (SHIMS 3.0)** |
|---|---|---|---:|
| P1 | Accurate | Incomplete | 28844 |
| P2 | Accurate | Accurate | 25534 |
| P3 | Incomplete | Missing | 36270 |
| P4 | Missing | Missing | 29842 |
| P5 | Missing | Missing | 105400 |
| P6 | Missing | Incomplete | 34928 |
| P7 | Accurate | Missing | 28530 |
| P8 | Missing | Incomplete | 53154 |
| P9 | Missing | Incomplete | 119578 |
| P10 | Missing | Accurate | 9184 |
| P11 | Incomplete | Incomplete | 160682 |
| P14 | Incomplete | Incomplete | 41640 |
| P15 | Incomplete | Missing | 90284 |
| P16 | Missing | Incomplete | 102504 |
| P17 | Accurate | Accurate | 14737 |
| P21 | Incomplete | Incomplete | 37360 |
| P22 | Incomplete | Accurate | 9934 |
| P23 | Missing | Incomplete | 38530 |
| P24 | Missing | Accurate | 11228 |
| P25 | Accurate | Missing | 35368 |
| P26 | Accurate | Accurate | 49024 |

**Supplemental Table S4:** Status of palindromes identified in SHIMS 3.0 in other rhesus macaque X assemblies. Missing: No palindrome present in non-SHIMS 3.0 assembly; Incomplete: Part of palindrome present in non-SHIMS 3.0 assembly; Accurate: Full palindrome present in non-SHIMS 3.0 assembly.

| Palindrome | Mmul_8.0.1 | Mmul_10 | Arm length (SHIMS 3.0) |
|---|---|---|---|
| P1 | Missing | Accurate | 14749 |
| P2 | Incomplete | Accurate | 42783 |
| P3 | Missing | Accurate | 35266 |
| P4 | Missing | Accurate | 11323 |
| P5 | Missing | Accurate | 19290 |
| P6 | Missing | Accurate | 15572 |
| P7 | Incomplete | Accurate | 24490 |
| P8 | Incomplete | Missing | 38496 |
| P9 | Incomplete | Incomplete | 81971 |
| P10 | Accurate | Accurate | 6574 |
| P11 | Incomplete | Missing | 106186 |
| P12 | Missing | Accurate | 12978 |
| P18 | Missing | Incomplete | 47531 |
| P19 | Missing | Incomplete | 14175 |
| P21 | Accurate | Accurate | 21963 |
| P24 | Missing | Accurate | 8398 |
| P25 | Missing | Missing | 103488 |
| P26 | Missing | Accurate | 46855 |

**Supplemental Table S5:** Conservation of X palindrome gene families. Values shown are the number of intact gene copies found in the given species. "Region" refers to the human region in which the gene family is found.

| Gene | Palindrome | Region | Human | Chimpanzee | Rhesus macaque |
|---|---|---|---|---|---|
| *CENPVL* | P2 | Arm | 2 | 2 | 2 |
| *MAGED4* | P3 | Arm | 2 | 2 | 2 |
| *FAM156* | P6 | Arm | 2 | 2 | 2 |
| *USP51* | P7 | Spacer | 1 | 1 | 1 |
| *CXorf49* | P8 | Arm | 2 | 2 | 2 |
| *DMRTC1* | P9 | Arm | 2 | 2 | 2 |
| *FAM236B* | P9 | Arm | 2 | 2 | 0 |
| *FAM236D* | P9 | Arm | 2 | 2 | 0 |
| *PABPC1L2* | P10 | Arm | 2 | 2 | 2 |
| *NXF2* | P11 | Arm | 2 | 2 | 2 |
| *TCP11X2* | P11 | Arm | 2 | 2 | 2 |
| *MAGEA12* | P21 | Spacer | 1 | 1 | 0 |
| *CSAG1* | P21 | Spacer | 1 | 0 | 0 |
| *MAGEA2* | P21 | Arm | 2 | 2 | 1 |
| *MAGEA3* | P21 | Arm | 2 | 2 | 3 |
| *CSAG2* | P21 | Arm | 2 | 3 | 2 |
| *FLNA* | P24 | Spacer | 1 | 1 | 1 |
| *EMD* | P24 | Spacer | 1 | 1 | 1 |
| *CTAG1* | P25 | Arm | 2 | 2 | 4 |
| *IKBKG* | P25 | Arm | 1 | 1 | 2 |
| *F8A* | P26 | Arm | 2 | 2 | 2 |
| *H2AB* | P26 | Arm | 2 | 2 | 2 |

**Supplemental Table S6:** Purifying selection on X palindrome genes. dN/dS values were calculated using the basic model in PAML (model=0,NSites=0).

2ΔL (observed vs. 1): Likelihood ratio test for observed dN/dS value versus null hypothesis (dN/dS = 1)

2ΔL (M1a vs. M2a): Likelihood ratio test for neutral evolution versus positive selection

| Gene | dN | dS | dN/dS | 2ΔL (observed vs. 1) | 2ΔL (M1a vs. M2a) |
|---|---|---|---|---|---|
| *PABPC1L2* | 0 | 0.0472 | 0.0001 | 16.94*** | 0 |
| *FLNA* | 0.0037 | 0.1483 | 0.02472 | 584.08*** | 0 |
| *IKBKG* | 0.0062 | 0.1403 | 0.04421 | 68.52*** | 0.18 |
| *F8A* | 0.0055 | 0.0832 | 0.06665 | 32.38*** | 0 |
| *MAGED4* | 0.0099 | 0.0695 | 0.14308 | 46.38*** | 0 |
| *CENPVL* | 0.0242 | 0.0998 | 0.24204 | 12.5*** | 0 |
| *FAM156* | 0.0202 | 0.0825 | 0.24445 | 10.56** | 0 |
| *USP51* | 0.0125 | 0.039 | 0.31986 | 12.16*** | 0.64 |
| *H2AB* | 0.066 | 0.1925 | 0.34267 | 6.70** | 0 |
| *EMD* | 0.0369 | 0.096 | 0.38423 | 7.68** | 4.18* |
| *NXF2* | 0.0334 | 0.0649 | 0.51501 | 7.28** | 0 |
| *MAGEA2* | 0.063 | 0.1078 | 0.58423 | 3.84 | 0 |
| *MAGEA3* | 0.1432 | 0.2091 | 0.68475 | 4.12* | 3.18 |
| *CXorf49* | 0.0725 | 0.1036 | 0.69933 | 3.06 | 0.86 |
| *CTAG1* | 0.1309 | 0.1751 | 0.74787 | 1.26 | 2.72 |
| *TCP11X* | 0.0406 | 0.0525 | 0.77242 | 0.6 | 0 |
| *DMRTC1* | 0.0165 | 0.0197 | 0.83446 | 0.08 | 0 |
| *CSAG2* | 0.0958 | 0.0866 | 1.10635 | 0.06 | 8.86** |

*p<0.05
**p<0.01
***p<0.001

**Supplemental Table S7:** Rates of P17 spacer deletions in azoospermic and oligozoospermic men

| Dataset | # men | # P17 deletions | % P17 deletions |
|---|---|---|---|
| 1000 Genomes | 944 | 126 | 13.3 |
| dbGaP phs001023 (control) | 292 | 54 | 18.5 |
| dbGaP phs001023 (azoospermia) | 286 | 47 | 16.5 |
| Oligozoospermia | 562 | 68 | 12.1 |

**Supplemental Table S8:** Chimpanzee clones used in this project that were previously sequenced and deposited in GenBank.

| Clone | Palindrome |
|---|---|
| CH251-26J9 | P1 |
| CH251-17J20 | P11 |
| CH251-98I1 | P12 |
| CH251-55N11 | P13 |
| CH251-52P5 | P21 |
| CH251-498N14 | P25 |
| CH251-25B20 | P26 |

**Supplemental Table S9:** PCR primers for gels shown in Supplemental Fig. 14.

| Feature | Forward primer (5' to 3') | Reverse primer (5' to 3') | Notes |
|---|---|---|---|
| P1 breakpoint | CCTCCTCCGTGTTTTTCTGA | CACAAGACAGGTGCAAGGAA | |
| P2 outer arm | CCCTCATCAAAAGGTAGGGG | CTGGGTAAGGAGATGGGGAT | |
| P2 spacer | CGTGCGTGTGTACCATCTTT | AGCTGACTTACATGGAGGGG | |
| P5 breakpoint | AGAAGGAGTCTCACTTTTGTCGCCCAAG | GCCTCCCAAAGTGTCTTTGTTCAGTTCA | Long range PCR |
| P8 breakpoint | AGACTGGGTGTTGCGAACAGACAAAAAC | GGATTTGTCTGAGAACTCATTCTTGGCG | Long-range PCR |
| P8 inner arm | TCCCACTGCTCTGCATCC | CTGGAAGAAGATCTTTATCCTGC | |
| P11 breakpoint | AATCCACAGGGGACAGCTC | TGTGGGGATAGGAAGTGACA | |
| P11 inner arm | GCAGGAGTTGCTTCTGTTACTG | TTTGAGTTTGGCTTTCCTGG | |
| P11 spacer | TCTGTTGAATATGCTCCACACC | TAGTGCAAATTGCTTTCCAGTC | |
| P17 breakpoint | TCAAAGTTGAAGGGTGTGGC | TTTGGCAATTCTTCCCTGTC | |
| P17 inner arm | AAAGCAAGCTCCTAAGGATGTG | GGCATCATCCAAACAAGTGG | |
| P17 outer arm | ATTCGAATGCTGACTCCCAC | GGGAGCTGAACTGCTGTACC | |
| P22 breakpoint | AGTACCACACAGAGAGGGAGC | GAGGTCAGGCAAGGAAAGAG | |
| P22 flanking | AACCATGGTCCCAAAATTCA | TCAGCAGTCAACCAGCATTC | |
| P22 spacer | TGACCATGACTGTGGGAGAA | CAGCCCCTGCTCAAGACTAC | |
| P25 breakpoint | TCATAGGCTGTTGATGACGG | CGTGATCCCCAAAGGTTG | |
| P25 inner arm | CACTGTGTCCGGCAACATAC | TCTGTTCTGAGACCCTGTGC | |
| P25 spacer | TCACACGCTGGTAATTGCAT | CAGCCCTCAGAAGAATTTGC | |

**Supplemental Table S10:** Chimpanzee and rhesus macaque clones sequenced for this project and deposited in GenBank

| Clone | Accession |
|---|---|
| CH250-106M20 | AC280444 |
| CH250-114J18 | AC280531 |
| CH250-119L11 | AC280481 |
| CH250-120L20 | AC280562 |
| CH250-136N6 | AC280430 |
| CH250-137I15 | AC280580 |
| CH250-138B21 | AC280536 |
| CH250-149O24 | AC280566 |
| CH250-150I6 | AC280452 |
| CH250-163K20 | AC280436 |
| CH250-168E3 | AC280520 |
| CH250-174F12 | AC280455 |
| CH250-184A21 | AC280508 |
| CH250-191K20 | AC280440 |
| CH250-197O3 | AC280457 |
| CH250-214O8 | AC280424 |
| CH250-228D11 | AC280538 |
| CH250-234D7 | AC280571 |
| CH250-236O7 | AC280541 |
| CH250-240H14 | AC280414 |
| CH250-257F3 | AC280575 |
| CH250-257M3 | AC280477 |
| CH250-25I12 | AC280437 |
| CH250-273C12 | AC280539 |
| CH250-280C5 | AC280451 |
| CH250-300J22 | AC280417 |
| CH250-312L23 | AC280517 |
| CH250-313D10 | AC280454 |
| CH250-318K15 | AC280486 |
| CH250-371L16 | AC280564 |
| CH250-396M7 | AC280543 |
| CH250-397P11 | AC280441 |
| CH250-398K19 | AC280504 |
| CH250-412K19 | AC280483 |
| CH250-417G7 | AC280442 |
| CH250-420A18 | AC280569 |
| CH250-424H13 | AC280467 |

| | |
|---|---|
| CH250-436M9 | AC280526 |
| CH250-440K2 | AC280563 |
| CH250-462M8 | AC280473 |
| CH250-486E21 | AC280527 |
| CH250-487N16 | AC280453 |
| CH250-491H11 | AC280464 |
| CH250-493M11 | AC280503 |
| CH250-498I16 | AC280468 |
| CH250-499B10 | AC280432 |
| CH250-503C21 | AC280489 |
| CH250-503N19 | AC280524 |
| CH250-504P11 | AC280429 |
| CH250-516N14 | AC280555 |
| CH250-530N5 | AC280476 |
| CH250-540J3 | AC280456 |
| CH250-541H5 | AC280425 |
| CH250-547J16 | AC280475 |
| CH250-563M7 | AC280492 |
| CH250-57C9 | AC280498 |
| CH250-80G22 | AC280568 |
| CH250-87B7 | AC280549 |
| CH250-92B13 | AC280534 |
| CH250-94G2 | AC280518 |
| CH250-95D17 | AC280449 |
| CH251-130O9 | AC280561 |
| CH251-160A4 | AC280458 |
| CH251-161L14 | AC280465 |
| CH251-172F20 | AC280557 |
| CH251-177B21 | AC280525 |
| CH251-183G21 | AC280578 |
| CH251-189G13 | AC280560 |
| CH251-239P10 | AC280533 |
| CH251-240O17 | AC280544 |
| CH251-261H21 | AC280545 |
| CH251-277H18 | AC280556 |
| CH251-285D14 | AC280499 |
| CH251-292E19 | AC280434 |
| CH251-316L7 | AC280500 |
| CH251-346A10 | AC280446 |
| CH251-34N14 | AC280480 |

| | |
|---|---|
| CH251-385I8 | AC280416 |
| CH251-389B7 | AC280426 |
| CH251-397P16 | AC280507 |
| CH251-4M24 | AC280567 |
| CH251-504H5 | AC280540 |
| CH251-506D4 | AC280415 |
| CH251-50L15 | AC280523 |
| CH251-514B7 | AC280488 |
| CH251-542A6 | AC280462 |
| CH251-542D16 | AC280579 |
| CH251-542E16 | AC280512 |
| CH251-550E20 | AC280495 |
| CH251-565G15 | AC280459 |
| CH251-571K4 | AC280521 |
| CH251-58J24 | AC280448 |
| CH251-635P13 | AC280558 |
| CH251-639F23 | AC280574 |
| CH251-64D22 | AC280469 |
| CH251-651H9 | AC280522 |
| CH251-654E24 | AC280553 |
| CH251-657L4 | AC280546 |
| CH251-658J15 | AC280445 |
| CH251-65E21 | AC280530 |
| CH251-671I19 | AC280576 |
| CH251-673E12 | AC280463 |
| CH251-677L24 | AC280565 |
| CH251-702N4 | AC280482 |
| CH251-737G9 | AC280491 |
| CH251-73C22 | AC280423 |
| CH251-83H5 | AC280548 |
